# Induction of cortical Par complex polarity by designed proteins causes cytoskeletal symmetry breaking in unpolarized mammalian cells

**DOI:** 10.1101/2022.03.15.484321

**Authors:** Joseph L. Watson, Ariel J. Ben-Sasson, Alice Bittleston, James D. Manton, David Baker, Emmanuel Derivery

## Abstract

Polarized cells rely on a polarized cytoskeleton for polarized trafficking, oriented migration and spindle orientation during asymmetric cell division. While cytoskeleton remodeling machineries have been extensively characterized at the molecular level, how polarity signaling at the cortex controls remodeling of the cytoskeleton in the cytosol remains elusive. In particular, how the Par complex, the conserved mastermind of polarity during asymmetric cell division, gets assembled and functions is not understood at the molecular level. Here, we dissected the logic of the Par complex pathway by capitalizing on designed proteins able to induce spontaneous symmetry breaking of the cortex in populations of naïve, unpolarized cells. We found that the primary kinetic barrier to Par complex assembly is the relief of Par6 autoinhibition, and that inducing Par complex cortical polarity was sufficient to induce two key hallmarks of asymmetric cell division in unpolarized cells: spindle orientation and central spindle asymmetry. These two outputs of the Par complex are separately controlled: spindle orientation is determined by Par3 and does not require the kinase activity of aPKC, while central spindle asymmetry solely depends on an asymmetric activity of aPKC at the cortex. Our work shows how polarity information flows between the cortex and the cytosol despite its diffusive nature, and paves the way towards induction of asymmetric cell division in cultured cells.

## Introduction

The Par complex is composed of three subunits, namely Par3, Par6 and the kinase aPKC, which are well conserved from flies to mammals. During asymmetric cell division, the Par complex forms cortical caps in metaphase, which are required to orient the mitotic spindle and to ensure the asymmetric segregation of cell fate determinants and thereby asymmetric cell fate determination between the daughter cells (Morin and Bellaïche, 2011). A conserved feature of asymmetric cell division is the polarized trafficking of signalling organelles to enhance the robustness of asymmetric cell fate determination (Coumailleau et al., 2009; Derivery et al., 2015; Katajisto et al., 2015; Kawaguchi et al., 2013; Kressmann et al., 2015; Loeffler et al., 2019; Montagne and Gonzalez-Gaitan, 2014; Zhao et al., 2020). There again, the Par complex has been proposed to orchestrate the necessary polarization of the cytoskeleton to support these polarized trafficking events. For instance, in flies, the Par complex is required for symmetry breaking of the anaphase midzone, also known as central spindle, which in turn biases the polarized trafficking of endosomes containing cell fate determinants towards one daughter cell (Derivery et al., 2015).

However, the assembly sequence of the Par complex is not understood at the molecular level. Indeed, while each subunit of the Par complex can bind to the two others, the triple complex is not always constitutively assembled (Morais-de-Sá et al., 2010; Soriano et al., 2016; see also Fig. S1A-C), and the way in which these proteins interact with one another to form a stable complex is a longstanding question in the field. For instance, aPKC is known to phosphorylate Par3 (Lin et al., 2000), but this inhibits the binding between aPKC and Par3 and therefore assembly of the complex. To allow complex formation, it has been proposed that Par6 binding could inhibit the kinase activity of aPKC (Atwood et al., 2007; Wirtz-Peitz et al., 2008; Yamanaka et al., 2001), although this is controversial (Graybill et al., 2012). This negative effect of aPKC phosphorylation on Par complex assembly is particularly enigmatic, since the way Par complex caps controls the cortical localization of fate determinants such as Numb is indeed through aPKC-mediated phosphorylation (Smith et al., 2007), so there must be a mechanism by which the assembled Par complex retains aPKC activity *in vivo*. Finally, while it is well established that the Par complex clusters at cell-cell junctions in polarized cells (Pickett et al., 2019), and that Par complex caps are formed of coalesced clusters (Kono et al., 2019), the specific contribution of clustering to Par complex assembly is not understood.

Similarly, the molecular mechanisms by which cortical caps of the Par complex control symmetry breaking of the microtubule cytoskeleton are also elusive. While the molecular cascade linking the Par complex to spindle orientation has been delineated in flies (Morin and Bellaïche, 2011), whether the same cascade holds true in mammals, and whether these proteins are sufficient to orient the spindle in the absence of other polarity pathways remains unclear. Furthermore, while it is known that the Par complex is upstream of spindle midzone asymmetry during asymmetric cell division in flies (Derivery et al., 2015), how this occurs at the molecular level is unknown. In this system, the central spindle displays an asymmetry in microtubule density, with one side having more microtubules than the other. This is a particularly enigmatic observation, as first, the Par complex is at the cortex and the spindle in the middle of the cell, so it is not clear how information flows between the two, and, second it is hard to envision that a gradient of microtubule regulators could exist in the cytosol of small cells given the highly diffusive nature of the cytosol.

*In vitro* reconstitution with purified components has consistently led to significant leaps in our molecular understanding of cellular processes, as it allows one to delineate what is *sufficient* for a given phenomenon to occur, rather than just what is *required*. By analogy, the *in vivo* reconstitution of polarity in otherwise unpolarized mammalian cells has been a long-standing goal of the field to increase our molecular understanding of cytoskeleton symmetry breaking and asymmetric cell division. Indeed, by artificially polarizing Par complex components during division, it would be possible to bypass feedback loops with other polarity pathways and the actin cortex, and thus to specifically interrogate which molecular interactions are *sufficient* for Par complex assembly and its downstream effects on the cytoskeleton. However, while pioneering progress has been made towards this goal, current methods are either restricted to fly cells (Johnston et al., 2009; Kono et al., 2019), or require external input via optogenetics in a microscope at the single cell level (Okumura et al., 2018), therefore precluding the systematic dissection of polarity pathways at the population level in mammalian cells.

Here, we capitalized on protein design to induce spontaneous long-term cortical polarity of virtually any protein of interest in otherwise unpolarized cells, and applied this novel assay to dissect the molecular logic of the Par complex in mammals. We found that inducing Par complex polarity was sufficient to induce three key hallmarks of asymmetric cell division in unpolarized cells, namely spindle orientation, asymmetric segregation of cortical components and central spindle asymmetry. Importantly, these events can be molecularly untangled: spindle orientation is determined by Par3 and does not require the kinase activity of aPKC, while central spindle asymmetry solely depends on an asymmetric activity of aPKC at the cortex. Furthermore, since our method allows the temporal separation of Par complex assembly from its downstream effects onto the cytoskeleton, this allowed us to quantitively and orthogonally study the assembly pathway of the Par complex. This revealed that the main kinetic barrier to Par complex assembly is the relief of Par6 autoinhibition, and that the kinase activity of aPKC has little quantitative effect on Par complex assembly *in vivo*. Our work thereby explains how Par complex caps can exhibit aPKC activity to induce their downstream effects on cytoskeleton symmetry breaking and the asymmetric segregation of fate determinants.

## Results

### Induction of cell polarity from the outside using designed protein arrays

To shed light on the molecular mechanisms underlying Par complex assembly and function during asymmetric cell division in mammals, we sought to induce acute cortical polarity of Par complex subunits in otherwise unpolarized cells. As a model system of unpolarized cells, we chose NIH/3T3 fibroblasts, in which the endogenous Par complex is unassembled with very little colocalization between Par3 and aPKC, both at the cortex and in the cytosol, except at inter-cell boundaries where the Par complex sometimes appears as punctate structures (Fig. S1A-C). In these unpolarized cells, the Par complex does not form caps during mitosis, and therefore the link between Par complex asymmetry and mitotic spindle is lost and the division is symmetric (in this paper, we consider asymmetric cell division as a division displaying physical asymmetries such as the spindle and cortical caps, rather than asymmetry in cell fate).

We first explored the possibility of inducing cell polarity using designed protein arrays. Our method relies on a synthetic transmembrane construct that is rapidly crystalized *from the outside* through assembly of a designed 2D protein material (Fig. 1). This material consists of two components, A(d) and B(c)-GFP, where B(c)-GFP binds to the transmembrane segment (via an anti GFP nanobody named GBP for “GFP Binding Peptide” (Kirchhofer et al., 2010), Fig. 1A), and A(d) clusters the B(c) component into a hexagonal array (Ben-Sasson et al., 2021) (Fig. 1A). Clustering with this method is fast (~20s Fig. 1B), efficient −~80 targets per diffraction limited spot (Ben-Sasson et al., 2021)- and occurs simultaneously and homogenously over the entire cell surface in interphase (Fig. 1C, Fig. S2A,B and Movie S1 for sub-cellular light sheet imaging). The resulting arrays are stable at the cell surface, as the material has been specifically engineered to evade endocytosis (Ben-Sasson et al., 2021), but we did not previously investigate what happens over long incubation periods (several hours). Remarkably, we found that these clusters spontaneously coalesce into cortical caps when cells round up during mitosis (Fig. 1D, Movie S2).

**Figure 1.**
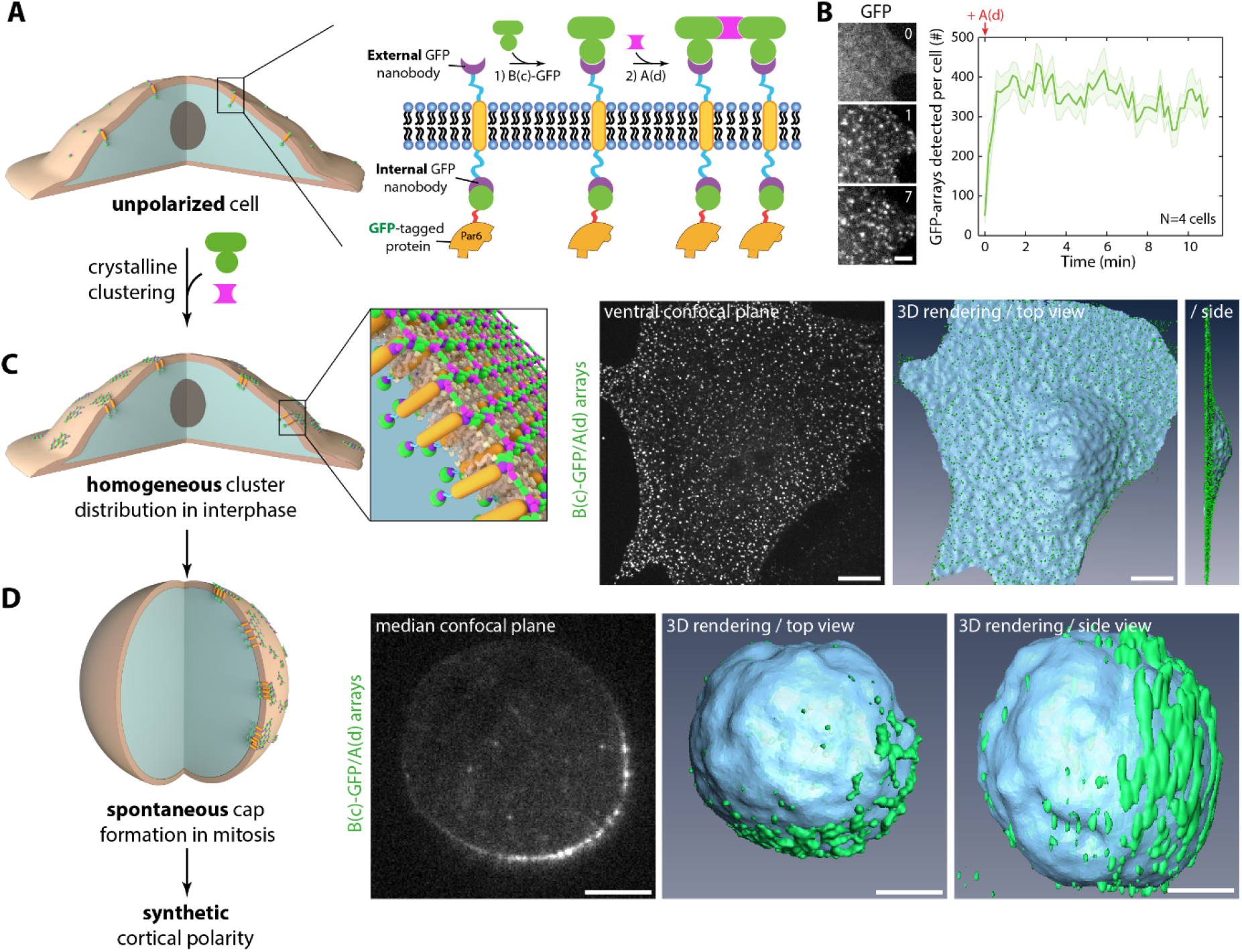
Artificial symmetry breaking of the cortex using protein design. (A) 3T3 cells stably coexpressing GBP-TM-GBP and a GFP-fused protein of interest were incubated with B(c)GFP then A(d) to induce rapid clustering. (B) Left: 3T3 cells stably coexpressing GBP-TM-GBP and GFP were incubated with B(c)GFP then A(d) and imaged by SDCM. Elapsed time in minutes. Right: Mean (+/-SEM) number of clusters per cell over time in cells. Clustering reaches steady state in ~20s. (C) Cells as in (B) were imaged 10 min after clustering by SDCM (middle panel: single confocal plane). Right panels show 3D views of the confocal stack shown in the left panel. Clusters, which correspond to hexagonal arrays of the protein of interest, are homogenously distributed at the cell cortex in interphase and thus allow the quantitative study of candidate protein recruitment to a clustered protein of interest (left panel). (D) Array assembly was triggered at the surface of 3T3 cells stably expressing GFP-LGN and GBP-TM-GBP as in (A), and cells were then stalled in mitosis for >12h with nocodazole before imaging by SDCM. Image corresponds to single confocal plane. Right: 3D rendering of confocal stack shown in the left panel. Note the partitioning of arrays into an asymmetric cortical cap during mitosis. Scale bars: 10 μm (C), 5 μm (D) and 2 μm (B).

The spontaneous coalescence of arrays into cortical caps was not due to the reagent used to stall cells in mitosis, as partitioning was evident in non-stalled cells (Fig. S3A-B), or even non-mitotic trypsinized cells (Fig. S3C). Similarly, this did not depend on the cell type (Fig. S3D, Movie S3), nor on the protein fused/relocalized to the transmembrane construct (Fig. 1D *versus* Fig. S3B,C). Lastly, cap formation is not specific to our transmembrane construct, as it occurs with native receptors, such as Notch, when functionalizing the clustering material with its ligand DLL4 (Fig. S3E,F). Rather, this partitioning seems driven by an energy-minimization mechanism. Using a membrane tension/lipid packing probe, Flipper-TR (Colom et al., 2018), we found that the Flipper-TR fluorescence lifetime was significantly increased in the membrane underlying the formed cap compared to the surrounding membrane (Fig. S3G,H and methods). This suggests that the 2D arrays exert some local change in membrane tension/lipid packing, creating a line tension, which in turn leads to cap formation as the cell would tend to minimize global energy (Fig. S3I). Alternatively, the clusters could have affinity for existing zones of high cortical tension. While the exact physics of cap formation are beyond the scope of this paper, the remarkable property of the clusters to spontaneously coalesce into caps when cells round up allows the reconstitution of the cortical symmetry breaking of virtually any protein of interest within an entire population of dividing cells.

### Par complex assembly is sequential and can occur via multiple pathways

We next took advantage of the protein arrays to investigate the molecular assembly pathway of the Par complex. Indeed, while each subunit of the Par complex can bind to the two others, the triple complex is not always assembled (Fig. S1A-C), and how the three subunits come together is unclear. To probe the various possible assembly pathways of the Par complex, and which molecular interaction are critical in each case, we systematically clustered each subunit (or mutants thereof) into arrays and quantitatively followed whether the two other subunits could be recruited to these clusters at the endogenous level (see Table 1 for list of stable cell lines). We performed this analysis in interphase, when clusters are sparse at the cortex rather than forming caps, so as to temporally separate Par complex assembly from its downstream effects. This capitalizes on the robustness of our clustering method, which allow us to make quantitative measurements on tens of thousands of virtually identical crystalline arrays in dozens of cells (see methods for the automated 3D colocalization with PSF fitting pipeline used).

To cluster Par complex subunits, which are cytosolic targets, we used a bicistronic system where stably expressed, GFP-tagged Par proteins are relocalized to an independently-expressed transmembrane construct via a second, internal anti-GFP nanobody (Fig. 1A). This indirect arrangement ensured comparable expression of the transmembrane construct between stable cell lines, and therefore comparable cluster size (Ben-Sasson et al., 2021), as opposed to direct fusion of Par complex proteins to the transmembrane construct (Fig. S2C-E). Note that these clusters are not three-dimensional aggregates or condensates, as here both the spacing and orientation of each protein of interest is defined by the crystal lattice of the material (Ben-Sasson et al., 2021) and is hence invariant to the internal target protein (inter-protein distance ~8 nm).

While, surprisingly, colocalization between Par6A and aPKC or Par3 was low in baseline conditions (2 minutes post-clustering), artificially clustering Par6A was sufficient to induce the recruitment of both aPKC and Par3 over 60 min (Fig. 2A-C and Fig. S4A-E for controls and Fig. S5B for split images). As the kinetics of Par6A clustering are virtually instantaneous compared to the slow kinetics of aPKC recruitment (~20 s versus 1 h), this suggested that aPKC and Par6 do not form a stable multiprotein complex in cells, but rather assemble upon clustering. Indeed, measuring the intensity of aPKC per Par6A cluster (rather than the binary colocalization), revealed an increase over time (Fig. S5C). This establishes that clustering, a commonly reported feature of Par complex biology, *per se* drives assembly of the core Par complex. To shed light on the alternative assembly routes, we also clustered aPKC (Fig. S6), or Par3 (Fig. S7), which both induced Par complex assembly, following similar kinetics as those arising from Par6A clustering (Fig. S6I, S7D and see also supplementary discussion). This establishes that, while the Par complex is unassembled in basal conditions, clustering of any of its core subunits can induce complex assembly with similar kinetics, even in interphase.

**Figure 2.**
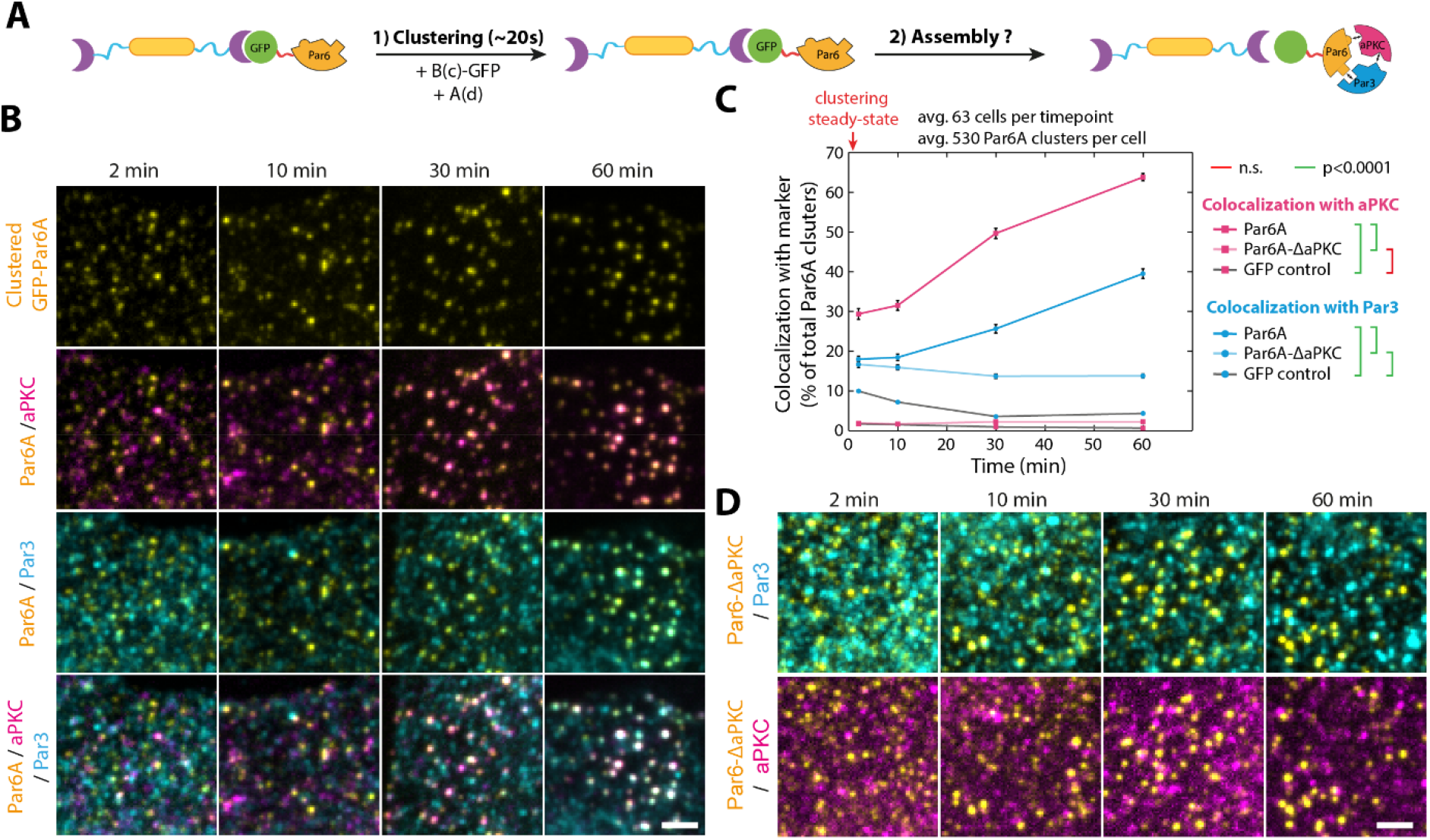
Sequential assembly of the core Par complex induced by clustering. (A) Principle of the experiment: GFP-fused Par6A, or mutant thereof, were clustered in 3T3 cells as in Fig. 1c and assembly of the Par complex was monitored. (B) GFP-Par6A was clustered as above, and the subsequent recruitment of endogenous aPKC and Par3 was monitored by immunofluorescence. For representation purposes, images were denoised with a “Wavelet a trous” filter (see methods). (**C**) Mean (+/-SEM) percentage of colocalization between clusters of indicated GFP-Par6A construct and endogenous aPKC/Par3 over time (see methods for details). Statistics: ANOVA2 using construct and timepoint as variables, followed by Tukey test (p value of each test indicated). Clustering of GFP alone, which does not bind to aPKC or Par3 is provided as a negative control to assess the accuracy of the method. aPKC gets gradually recruited to Par6A clusters, followed by Par3. (D) Clustering of GFP-Par6A^ΔaPKC^ does not trigger Par complex assembly. Note that throughout this paper, all transgenes and combination of transgenes are expressed from the same locus (see methods). Fig. S5 shows split-channel images of all multicolor panels presented in this figure. Scale bar: 2 μm.

Interestingly, aPKC recruitment to Par6A clusters preceded that of Par3 (Fig. 2B,C), suggesting a sequential assembly pathway where aPKC binding to Par6A might be a prerequisite for Par3 recruitment. To test whether aPKC binding was indeed required for Par3 recruitment to Par6, we clustered Par6A^ΔaPKC^, an established Par6 point mutant incapable of binding to aPKC (Hirano et al., 2005). Clustered Par6A^ΔaPKC^ was unable to recruit either aPKC or Par3 (Fig. 2C,D and Fig. S5B for split images). Therefore, aPKC binding to Par6 is required for full Par complex assembly downstream of clustered Par6. Interestingly, the different isoforms of Par6 were not identical in their ability to induce Par complex assembly upon clustering, with Par6A being the most and Par6B the least efficient (Fig. S5D-F).

Since Par3 binding to clustered Par6 first required aPKC recruitment, we wondered whether aPKC could recruit Par3 in the absence of Par6. To this end, we used aPKC^ΔPar6^, an established mutation abolishing Par6 binding (Hirano et al., 2005). aPKC^ΔPar6^ was not able to recruit Par3 when clustered, demonstrating that both aPKC and Par6 are required for Par3 recruitment to the Par complex (Fig. S6F,I). While tools to detect endogenous Par6 isoforms at these aPKC clusters were not available, we confirmed by mild transient overexpression that all three of the mammalian Par6 isoforms could be recruited into aPKC clusters to assemble the full aPKC/Par6/Par3 complex (Fig. S8). These results demonstrate that the interaction between Par6 and aPKC is absolutely required for quantitative recruitment of Par3 and assembly of the full Par complex.

### Relieving Par6 autoinhibition is the rate-limiting step in Par complex assembly

A tantalizing hypothesis to explain the delayed recruitment of aPKC and Par3 to Par6 clusters is that complex assembly depends on phosphorylation, as aPKC-mediated phosphorylation of Par3 reduces aPKC-Par3 binding but has been proposed to be inhibited by Par6 (Atwood et al., 2007; Lin et al., 2000; Wirtz-Peitz et al., 2008; Yamanaka et al., 2001) (Fig. S6A). We thus wondered whether these previous findings could explain our observations: Par6 and aPKC would bind to one another upon clustering, leading to the inhibition of aPKC and hence the subsequent recruitment of dephosphorylated Par3 (dephosphorylated due to the inhibited kinase activity of aPKC). A direct prediction of this paradigm is that clusters of a kinase-active aPKC mutant would not be able to recruit Par3 (as Par3 would be constitutively phosphorylated), and, conversely, clusters of a kinase-dead mutant of aPKC would be able to recruit Par3 in the absence of binding to Par6 (if the role of Par6 was simply to inhibit the kinase activity of aPKC). To our surprise, both established kinase-active (aPKC^active^) and kinase-dead (aPKC^dead^) mutants of aPKC recruited Par3 to the same extent as the wild-type (Fig. S6C-E,I), and combining the kinase-dead mutation and the Par6-binding mutant (i.e. aPKC^dead ΔPar6^) completely abolished Par3 recruitment and thus Par complex assembly (Fig. S6H,I). This implies that Par6 binding to aPKC is required for Par3 recruitment, and is not simply acting to inhibit the kinase activity of aPKC. Furthermore, this suggests that Par3 can be recruited to aPKC clusters even when phosphorylated (most likely indirectly via Par6). This was confirmed by clustering of Par3 phosphomimic mutants for the aPKC target sites, which only marginally affected assembly in the presence of Par6 (Fig. S7E and supplemental discussion for further details). This suggests that the kinase activity of aPKC is mostly dispensable for Par complex assembly and cannot explain the observed delay in this process.

So why is aPKC and Par3 recruitment to clustered Par6 so slow? Another hypothesis could be Par6 autoinhibition: aPKC binding could be required to ‘open’ Par6 to expose its Par3 binding domain. To test this possibility, we generated fragments containing the N-terminal (residues 1-121, henceforth Par6A^Nter^, which harbors the aPKC binding site) or C-terminal (residues 121-346, which harbors the Par3 binding site henceforth Par6^Cter^) halves of Par6 (Fig. 3A-B). Remarkably, clustering Par6A^Nter^ was able to recruit aPKC, but not Par3, whereas clustering Par6A^Cter^ was able to recruit Par3, but not aPKC (Fig. 3C-E, see also Fig. S9B for split channel images). Consistent with this, aPKC recruitment by Par6A^Nter^ was faster than Par6A full length (Fig. 3E). This was expected if the slow aPKC recruitment was the result of the need to relieve Par6 autoinhibition. Last, Par6A^Cter^ could recruit Par3, while the full length Par6A^ΔaPKC^ mutant could not, suggesting that full length Par6A indeed exists in an autoinhibited state (Fig. 3D).

**Figure 3.**
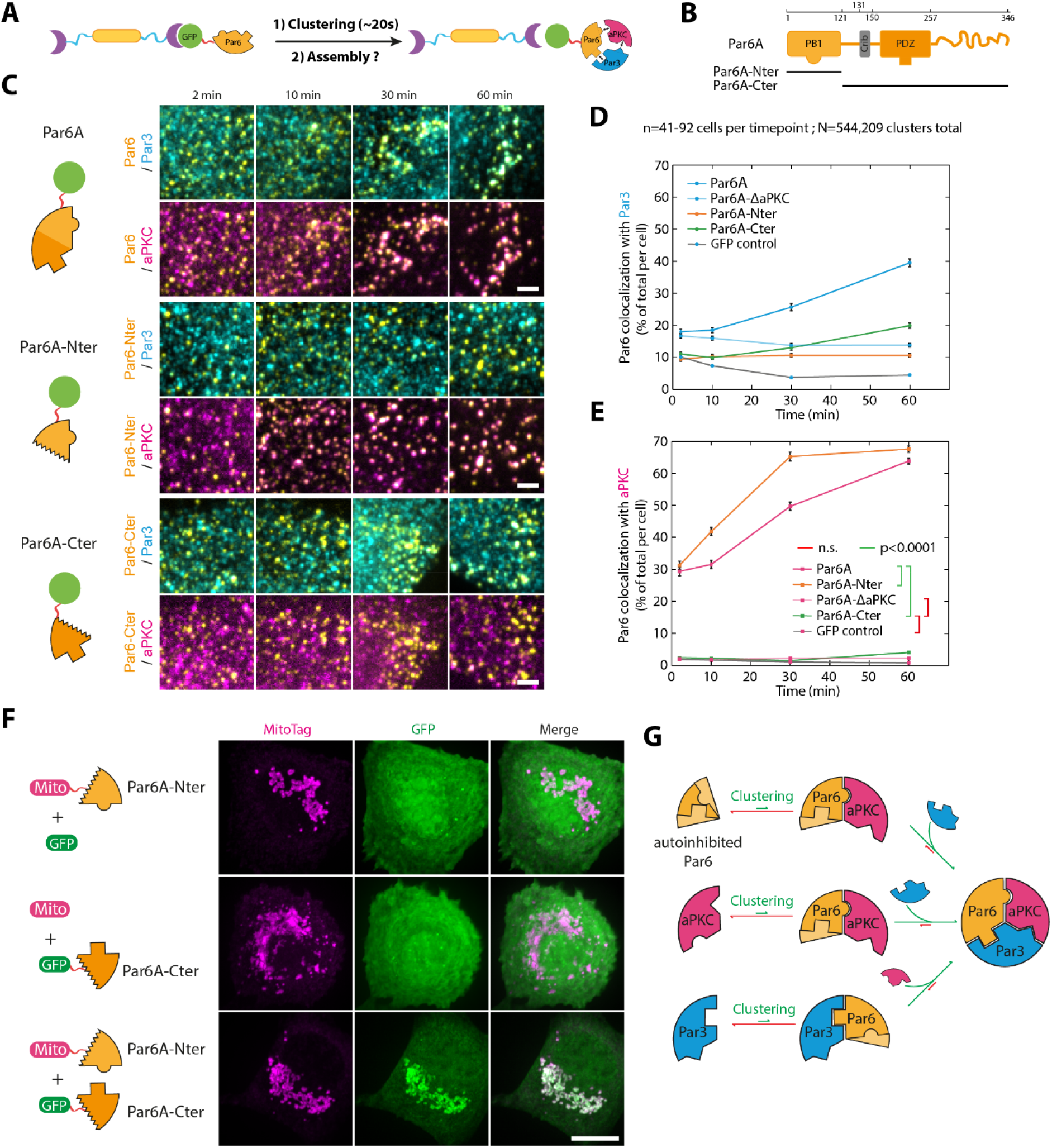
Relieving Par6 autoinhibition is the rate-limiting step in Par complex assembly. (**A**) Principle of the experiment: GFP-fused Par6A, or mutant thereof, were clustered in 3T3 cells as in Fig. 1c and assembly of the Par complex was monitored. (**B**) Domain organization of Par6A, with the N-terminal PB1 domain binding to aPKC and the C-terminal PDZ domain binding to Par3. (**C**) GFP-fused Par6A, or fragments thereof, were clustered in 3T3 cells and the recruitment of endogenous aPKC/Par3 was monitored by immunofluorescence (see also Fig. S9b for split-channel images). Images correspond to maximum intensity projection and images were denoised with a “Wavelet a trous” filter. (**D**,**E**) Mean (+/-SEM) percentage of colocalization between GFP-Par6A clusters and Par3 (**D**) or aPKC (**E**) over time. Statistics for (**D**): ANOVA to test interaction with time: Par6A: p<0.0001; Par6A^Cter^: p<0.0001; Par6A^Nter^: n.s.; Par6A^ΔaPKC^: p<0.01). Statistics for (**E**): ANOVA2 using construct and timepoint as variables followed by (p-value of each test indicated). Note that the curves for Par6A and the GFP control are the same as in Fig. 2, reproduced here for convenience. (**F**) The N-terminal domain of Par6A can bind to the C-terminus. 3T3 cells expressing GFP-Par6A^Cter^, and Par6A^Nter^ tethered to the mitochondria, or relevant controls, were imaged by SDCM. Images correspond to maximum intensity z-projection over the entire cell. Par6A^Nter^ can recruit Par6A^Cter^ to mitochondria, suggesting that the protein can fold on itself (see also Extended Data Figure 9c-d for reverse experiment and narrowing down of the interaction region). (**G**) Preferred route for assembly of the core Par complex as a function of the clustered subunit (see also supplementary discussion and Extended Data Figs. 5-9). Scale bars: 2 μm (**A**) and 10 μm (**D**).

This autoinhibitory binding between the two halves of Par6 was confirmed by relocalization experiments where mitochondria-targeted Par6A^Nter^ robustly recruited cytosolic GFP-Par6^Cter^ to mitochondria (Fig. 3F, see also Fig. S9C for narrowing down of the interaction region between the PB1 and PDZ domains). Likewise, mitochondria-targeted Par6^Cter^ robustly recruited GFP-Par6^Nter^ there (Fig. S9D). This interaction is likely direct, as when the two halves of Par6 were separately expressed in *E. coli*, where homologues able to recapitulate indirect binding are unlikely to exist, GST-Par6^Nter^ could specifically pulldown PC-Par6^Cter^ (Fig. S9E,F).

Integrating all these data, we propose the following paradigm for Par complex assembly (Fig. 3G, see also supplementary discussion for rationale of each branch). Our model has four key features: i) clustering of any of the core subunits can trigger assembly of the complex; ii) no matter the route, assembly is slow and regulated, the rate-limiting step being the ‘opening’ of an autoinhibited Par6; iii) there is a preferred assembly route depending on the subunit clustered; iv) the direct binding between aPKC and Par3 is not required for Par complex assembly, and the inhibitory effect of Par3 phosphorylation (via aPKC) on Par complex assembly does not quantitatively matter in the presence of Par6. This model rationalizes the sequential assembly observed in Fig. 2C: by increasing the local concentration of Par6 through clustering, aPKC is able to bind and open Par6 by outcompeting C-terminal binding to the N-terminus, and thereby allowing Par6 to bind to Par3. Similarly, our finding that the rate-limiting step in the assembly of the Par complex is the ‘opening’ of Par6 can also harmonize multiple discrepancies previously found in the literature (see discussion and supplemental discussion).

### An asymmetric cortex of the Par complex is sufficient to induce spindle orientation

Having rationalized the assembly pathways of the Par complex, we next wondered if the route used for assembly affects its outputs when forming a cap during mitosis. We first investigated whether the caps formed by our 2D arrays had any intrinsic effect on the orientation of division. Importantly, generating asymmetric caps of GFP had no effect on the orientation of division (Fig. 4A, Movie S4), with the angle of division being uncorrelated with either the position or the size/intensity of these control caps (Fig. 4B and Fig. S10A-B for individual datapoints). In stark contrast, targeting of any of the core Par complex subunits to the cap was able to robustly orient division to a similar extent (Fig. 4C-E, Movie S5-7). Importantly, an asymmetric cap of aPKC^dead^ (which, as aforementioned, can efficiently assemble the full Par complex) was as potent as its wildtype counterpart at inducing spindle orientation, indicating that the kinase activity of aPKC is not required for this process (Fig. 4E-F, Movie S8). Remarkably, asymmetric caps of the Par complex were also sufficient to induce another defining feature of asymmetric cell division, namely the preferential inheritance of polar caps by only one daughter cell (Fig. 4C-F, Movie S5-8). These results establish that an asymmetric cortex of the Par complex is sufficient to induce spindle orientation in unpolarized mammalian cells, and that this does not depend on the pathway used to assemble the Par complex, nor on the kinase activity of aPKC.

**Figure 4.**
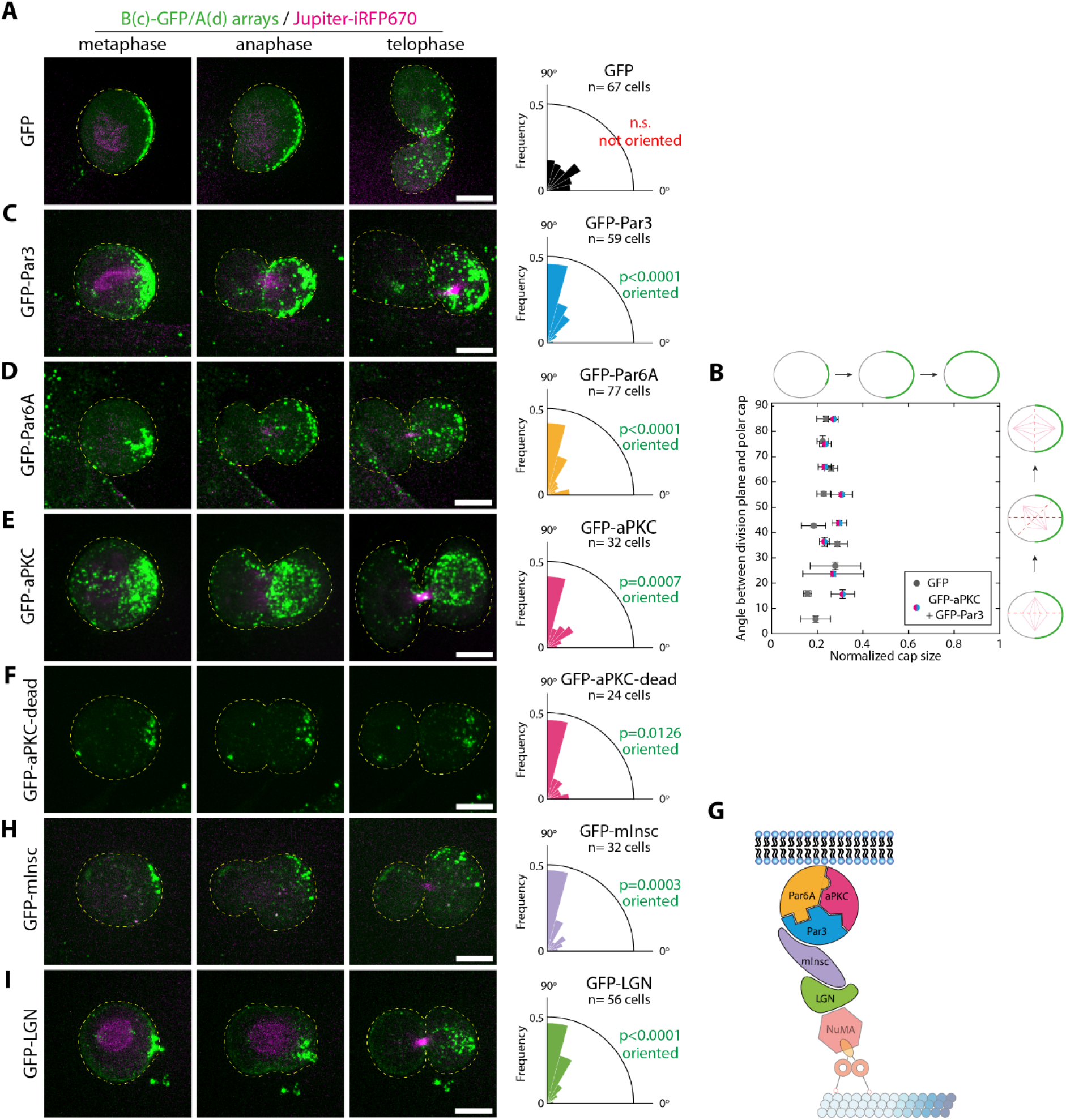
An asymmetric Par complex cortex is sufficient to orient the mitotic spindle. (**A,C-F**) Left: Array assembly was triggered at the surface of 3T3 cells stably co-expressing Jupiter-iRFP670, GBP-TM-GBP as well as indicated GFP-fusions. Cells were stalled in mitosis for >12h then imaged by SDCM upon release of the nocodazole block. In these conditions, array coalesce into asymmetric caps (Fig. 1d). Right: Angle between the division plane and the array cap as a function of the protein targeted to the cap. A 90° angle between division plane and cap corresponds to perfect orientation. Statistics: Unpaired Wilcoxon signed-rank test considering a hypothetical value of 45° (n: number of cells analysed). While the GFP control cells show random orientation, polar targeting of any of the core Par complex components (**B**: Par3, **C**: Par6, **D**: aPKC) or the kinase dead version of aPKC (**F**) induces spindle orientation and asymmetric segregation of the polar cap into one daughter cell in otherwise unpolarized cells. (**B**) Orientation of the division (angle between the division plane and the cap) as a function of the cap size. To have a normalized measurement between cells, cap size was expressed as a fraction of the cell perimeter, and was determined in metaphase in the equatorial plane. Note that since aPKC and Par3 cells have too few cells failing to orient their spindle, we pooled datasets to generate the plot across the whole range of orientation angles. (**G**) Cartoon illustrates the established (simplified) spindle orientation pathway (Morin and Bellaïche, 2011). (**H-I**) Asymmetric caps of indicated GFP fusions in dividing cells was induced and imaged as in **A.**mInsc (**H**) and LGN (**I**) caps are sufficient to induce spindle orientation and asymmetric cap segregation. Scale bars: 5 μm.

Remarkably, we found no correlation between the cap size and the robustness of spindle orientation upon targeting of core Par complex components to the cap (Fig. 4B and S10A,C for individual datapoints). This suggests that the molecular composition of the cap, rather than its geometry, is the determining factor for spindle orientation. We thus wondered if asymmetric caps of other proteins previously shown to be required for spindle orientation (Fig. 4G) were also sufficient to do so in unpolarized mammalian cells. In particular, caps of Inscutable (mInsc) and LGN were found to robustly orient the spindle along the established polarity axis, and thereby led to asymmetric inheritance of the polar cap (Fig. 4H,I; Movie S9,10).

### Asymmetric, cortical aPKC activity controls central spindle symmetry breaking

Finally, we wondered if an asymmetric cortex of the Par complex was sufficient to induce central spindle symmetry breaking. Indeed, we previously showed that during the asymmetric division of *Drosophila* Sensory Organ Precursors (SOP), the anaphase-specific spindle, known as the central spindle, also undergoes symmetry breaking, whereby the anterior side of the central spindle contains more microtubules than the other (Fig. 5A and also ref (Derivery et al., 2015)). This asymmetric central spindle in turn orchestrates the polarized trafficking of signaling endosomes containing Notch and its ligand Delta towards only one daughter cell, thereby substantially contributing to asymmetric cell fate determination (Derivery et al., 2015). Yet, how this central spindle asymmetry arises, and if it is conserved in mammals, is unknown.

**Figure 5.**
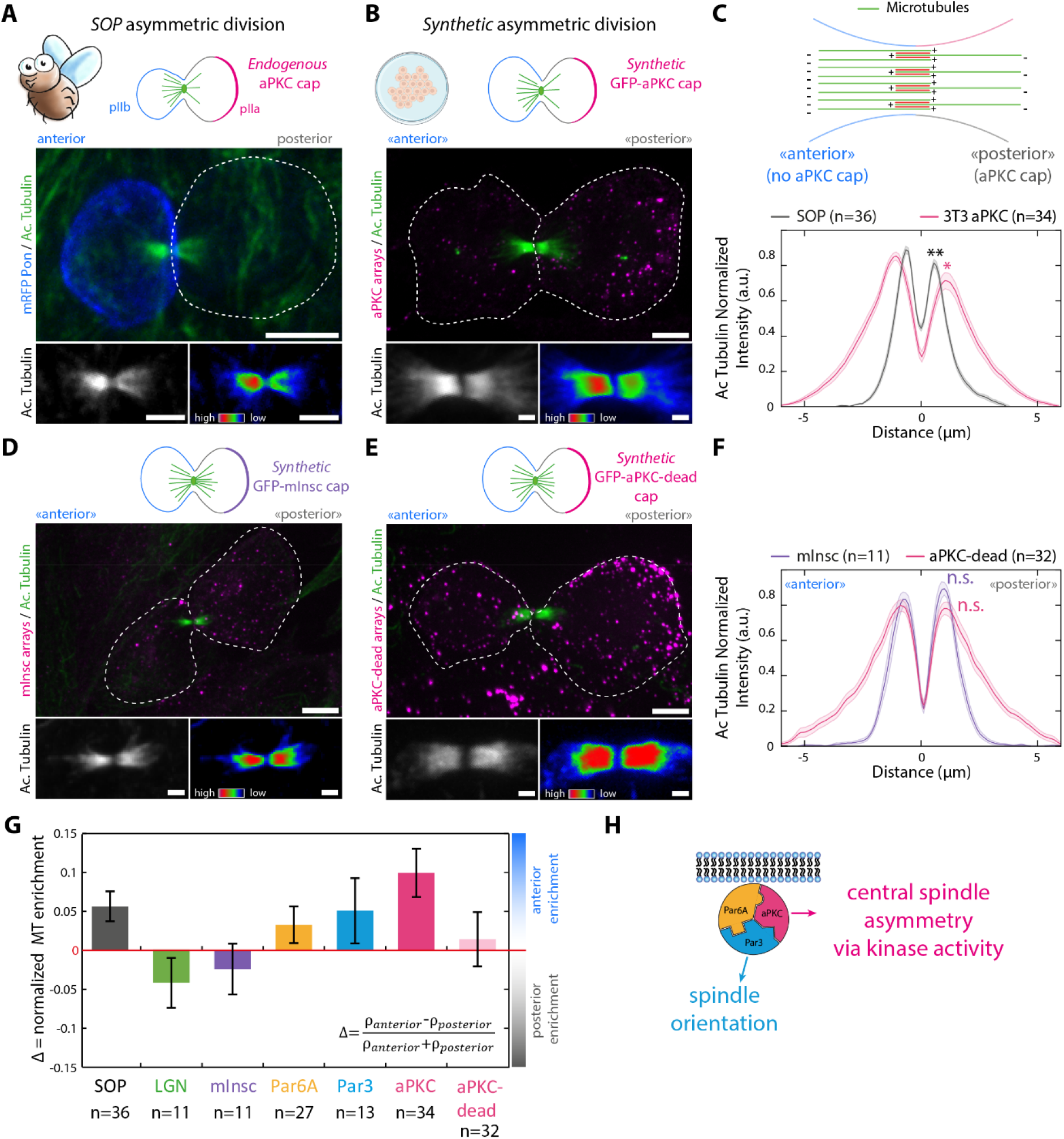
Asymmetric cortical aPKC activity controls central spindle symmetry breaking. (**A**) Drosophila Sensory Organ Precursor (SOP) cells expressing mRFP-Pon as a marker of the anterior pIIb cell (that is the cell not inheriting the core par complex cap) were fixed and immunostained for acetylated tubulin. Bottom panels: split-acetyl-tubulin channel with a grayscale (left) or Rainbow (right) lookup table. Note that the anterior side of the central spindle has a higher density of microtubules than the posterior. (**B**) Caps of array was triggered at the surface of 3T3 cells stably expressing GFP-aPKC and GBP-TM-GBP by stalling cells in mitosis for 12h. Nocodazole block was then released and cells were fixed after 30 min, before processing for acetyl-tubulin immunostaining. Note that the side of the central spindle in the cell that did not inherit the aPKC cap (“anterior”) has a higher density of microtubules than the “posterior” side, as in SOP cells. (**c**) top panel: proposed topology of the asymmetric central spindle adapted from ref (Derivery et al., 2015). Bottom panel: Acetyl-tubulin intensity pseudolinescan through the central spindle (mean+/-SEM; n: number of cells measured see methods). Statistics: paired t-test between the respective peak values in each cell (p=0.0068 for SOP and p=0.0107 for 3T3 aPKC). (**D-F**) 3T3 cells stably expressing GBP-TM-GBP and GFP-mInsc (**D**) or GFP-aPKC^dead^ (**E**) were processed and imaged as in (**B**), and central spindle asymmetry analyzed as in (**C**). Statistics: p=0.3232 for mInsc and p=0.7665 for aPKC kinase dead. (**G**) Normalized enrichment of microtubule density on the anterior side of the central spindle as a function of the protein targeted to the cap. The SOP central spindle is provided as a reference. While asymmetric caps of core Par complex components leads to central spindle symmetry breaking, this is not the case for LGN and mInsc or the kinase dead mutant of aPKC. (**H**) Summary of the findings presented in Figs. 4 and 5: the Par complex has two discrete outputs, spindle orientation, which goes through Par3 and does not rely on the kinase activity of aPKC, and central spindle asymmetry, which depends on aPKC phosphorylation of cytoskeleton targets. Images in this figure correspond to maximum intensity projections of the entire cell, or central spindle. Scale bars: 5 μm (**A**,**B**,**D**,**E**, top panels) and 1 μm (**A**,**B**,**D**,**E**, bottom panels).

Remarkably, upon reconstitution of an asymmetric cap of aPKC in unpolarized 3T3 cells, the central spindle also became asymmetric, in a very similar fashion to fly SOPs (Fig. 5B, see also C for average measurement of the microtubule density across the spindle). Note that the orientation is also conserved: the side that inherits the Par complex cap containing aPKC is the one that displays a lower microtubule density (Fig. 5A-C). This establishes that an asymmetric Par complex is sufficient to induce central spindle asymmetry in mammalian cells. Strikingly, while central spindle symmetry breaking also occurred upon targeting of the other core Par complex components to the cap (Par3/Par6), albeit to a lesser extent, it did not occur upon targeting of more downstream members of the spindle orientation pathway, such as mInsc and LGN (see Fig. 5D,F,G for mInsc and also Fig. S11A-C for Par3/Par6/LGN). This implies that central spindle asymmetry does not occur downstream of spindle orientation, or, in other words, that central spindle asymmetry and spindle orientation are two discrete outputs of Par complex activity that can be untangled (Fig. 5H).

The fact that mInsc caps do not induce central spindle asymmetry is surprising, since mInsc binds to both Par3 and LGN (Culurgioni et al., 2018), at least when clustered. One would therefore expect that mInsc clustering should not only induce spindle orientation (via LGN clustering), but also central spindle asymmetry via Par complex assembly (through Par3 clustering, see Fig. 4G). This provided a unique opportunity to probe the molecular details of the central spindle asymmetry pathway. Strikingly, while mInsc clusters recruited endogenous Par3, these mInsc-Par3 clusters did not recruit aPKC (Fig. S11D,E). Given our new framework for Par complex assembly (Fig. 3G), one reason for this could be that the indirect clustering of Par3 via mInsc is not efficient at “opening” Par6 in order to quantitatively recruit aPKC. This finding pointed towards aPKC being the key effector for central spindle asymmetry, which was in line with aPKC caps being more potent at inducing central spindle asymmetry than Par3/Par6 caps (Fig. 5G). This was confirmed by the fact that the kinase-dead mutant of aPKC (aPKC^dead^) while competent to robustly induce Par complex assembly and spindle orientation (Figs. S6, 4F), was reduced in its ability to induce central spindle symmetry breaking (Fig. 5E-G). This suggests that central spindle symmetry breaking is controlled by an asymmetric cortex of aPKC activity.

## Discussion

### Synthetic dissection of the input/output logic of polarity pathways

Here, we engineered a population of mammalian cells that spontaneously polarize a given protein of interest at their cortex just prior to division. Importantly, since cap formation with our method is an intrinsic property of our *de novo* designed polymer, our method can be adapted to virtually any other cell types/species (as we did in Fig. S3). Thus, our work generalizes previous pioneering work on the reconstitution of cortical polarity, which was either limited to fly cells (Johnston et al., 2009; Kono et al., 2019), or required external input via optogenetics (Okumura et al., 2018) and was thus restricted to single cells. Furthermore, a key advantage of our method is that since cortical targeting of the protein of interest is indirect via a nanobody, it allows independent control of the expression of the transmembrane segment, which controls the size of the clusters (Ben-Sasson et al., 2021), and that of the protein of interest, which is important for the stoichiometry of multiprotein complexes (Derivery and Gautreau, 2010)(Fig. S2).

Another major advantage of our method is that by design, it not only allows one to dissect the downstream effects of polarity proteins once assembled into a cap (i.e. the pathway outputs), but it also allows the study of the assembly of polarity signalling networks at the cortex (i.e. its inputs), all within the same experiment. Indeed, the inducible clustering of Par complex subunits in interphase, were no cap is formed, allowed us to systematically and quantitatively dissect the molecular assembly routes of the Par complex (Figs. 2–3), while clustering during mitosis allowed us the study of the effects of a *preassembled* Par complex cap on cytoskeleton symmetry breaking (Figs. 4–5).

It must be emphasized that the reconstitution of cortical polarity in cultured cells *de facto* enables live cell imaging at high spatio-temporal resolution, as we have done here. This is a key advantage over tissues and organoids, which have until now been required to study Par complex polarity in mammals, but where quantitative imaging is more challenging because of scattering. We envision that this key advantage of our method will be harnessed in the future to provide unprecedented details of the dynamics of symmetry breaking using lattice light sheet microscopy (Pamula et al., 2019) or into the structure of polarized cytoskeleton networks using Cryo Electron Tomography (Foster et al., 2022), both methods being readily applicable to cultured cells. Finally, given the generality of our method, we anticipate that it will prove useful to decipher the input/output logic of other polarity pathways beyond the Par complex, for instance at the immune synapse.

### A unified framework for Par-complex assembly

Integrating data from millions of arrays, we could systematically and quantitatively dissect the assembly sequence of the Par complex pathway and propose a unified paradigm for this process (Fig. 3G). Importantly, our results are highly consistent with previous work. In particular, we have confirmed that: i) Par6 and aPKC can interact, with aPKC^E85/R91^ and Par6^K19^ vital for this interaction (Hirano et al., 2005); ii) Par6 and Par3 can interact, with Par3^G600,602^ and Par6^121-257^ vital for this interaction (Liu et al., 2020); iii) Clustering/oligomerization of Par3 is required for Par complex assembly (Lin et al., 2000) and iv) The Par complex can assemble in clusters at cell-cell junctions (Pickett et al., 2019).

Our work, however, significantly extends these previous findings and provides key and novel insights into Par complex biology. In particular, we found that: i) The Par complex is mostly unassembled in single cells, with Par complex components largely cytoplasmic (Fig. 2); ii) Clustering of any core Par complex component can induce assembly of the tripartite complex (Figs. 2,S6,S7); iii) Par6 exists in an autoinhibited state, with the N- and C-termini interacting to prevent binding to aPKC and Par3 respectively (Fig. 3); iv) Clustering of any of the three components can relieve the autoinhibition of Par6 to allow binding to aPKC and Par3, but Par6 is most readily ‘opened’ by aPKC (Figs. 2,S6,S7); v) The role of the oligomerization domain in Par3 for Par complex assembly is solely to cluster Par3 and to allow the ‘opening’ of Par6 (Fig. S7).

Furthermore, our work also reconciles long-standing discrepancies in the field. In particular, while we confirmed that full length Par3 and aPKC can interact, with phosphorylation of Par3 by aPKC mildly inhibiting this interaction (Lin et al., 2000), we found that the kinase activity of aPKC is largely dispensable for Par complex assembly *in vivo*. Indeed, when Par3 is phosphorylated, the Par complex can still quantitatively assemble via Par6, without the direct aPKC-Par3 interaction (Figs. S6,S7). Therefore, our results rationalize how the Par complex can simultaneously be assembled and exhibit aPKC activity, and therefore how Par complex caps control the cortical distribution of fate determinants such as Numb, or, as we show here, central spindle symmetry breaking (see also supplementary discussion for further discussion of our results in context of the literature).

### Par complex polarity has two discrete effects onto cytoskeleton symmetry breaking

A major finding of this work is the demonstration that an asymmetric cortex of the Par complex has two discrete, independent outputs during mitosis: spindle orientation, which acts via Par3/mInsc/LGN but does not rely on the kinase activity of aPKC (Fig. 4), and central spindle asymmetry, which solely depends on the kinase activity of aPKC (Fig. 5). This key insight that the Par complex can independently affect two properties of the spindle (rotation and density) rationalizes why, in flies, some loss of function mutants were found to affect central spindle asymmetry without affecting spindle orientation (Derivery et al., 2015). This also demonstrates that central spindle asymmetry is not a fly oddity, but a conserved hallmark of asymmetric division.

Furthermore, our data also provide a new paradigm to explain the biophysics of central spindle symmetry breaking. Indeed, while it has been hard to envision that a gradient of cytosolic microtubule regulators could exist in cells due to the highly diffusive nature of the cytosol, our finding that central spindle asymmetry is controlled by cortical aPKC activity rather suggests the existence of an activity gradient. Indeed, if the activity of the microtubule regulator(s) responsible for central spindle asymmetry was regulated by aPKC-mediated phosphorylation, then an asymmetric aPKC cortex could lead to a steady state gradient of the phosphorylated regulator in the presence of cytosolic phosphatases. Hence, an activity gradient could exist despite diffusion. A fascinating question for future research is to test this paradigm, and to identify the aPKC target(s) responsible for how information flows between the cortex and the diffusive cytosol. It would also be interesting to know if this represents a general mechanism by which the Par complex imprints polarity onto the cytoskeleton of polarized cells in interphase, such as in epithelial cells. The assay described here should, in future, allow the direct interrogation of these questions.

Last, it is remarkable that naïve, unpolarized fibroblasts still express all the proteins needed to polarize their cytoskeleton provided the polarity of only one subunit of the Par complex is restored. This may suggest that all unpolarized cells are missing to divide asymmetrically is Par complex polarity. While beyond the scope of this study, it would be interesting to know if the reconstitution of Par complex caps is sufficient to induce other aspects of asymmetric cell division in fibroblasts, such as asymmetric segregation of cell fate determinants (Morin and Bellaïche, 2011), asymmetric gene expression, or polarized trafficking of endosomes (Daeden and Gonzalez-Gaitan, 2018), mitochondria (Katajisto et al., 2015) and/or lysosomes (Loeffler et al., 2019). Furthermore, our method could also be used to investigate the molecular basis of another hallmark of asymmetric cell division, namely cell size asymmetry. In particular the role of asymmetric cortical tension in this process (Delgado and Cabernard, 2020) could be decipher by inducing artificial caps of contractile actin with our method, or, similarly, our polymer could be combined with adhesive micropatterning to study how cell shape and cortical polarity are integrated to generate the final size of the daughter cells (Godard et al., 2021).

## Conclusion

In conclusion, this work highlights the power of synthetic biology and protein design to address cell biology questions in an orthogonal manner. Here, we used a crystalline polymer engineered to cluster specific proteins at a controlled density. This allowed us to uncouple clustering of the Par complex from oligomerization of Par3, which is presumably the driver of clustering in polarized cells (Liu et al., 2020). Indeed, by directly clustering Par6, rather than relying on indirect clustering through Par3, we were able to uncover a previously unappreciated feature of Par complex biology, namely Par6 autoinhibition as a kinetic barrier. Further, the key property of our polymer to spontaneously induce symmetry breaking of the cortex allowed us to demonstrate that an asymmetric cap of the Par complex is sufficient to induce hallmarks of asymmetric cell division such as spindle rotation and central spindle asymmetry. To our knowledge, this work represents the most complete *synthetic* reconstitution of asymmetric cell division to date, and paves the way for further delineation of the intracellular molecular mechanisms governing polarity and asymmetric cell division.

## Acknowledgments

This work has been supported by the Medical Research Council (MC_UP_1201/13 to E.D), the Human Frontier Science Program (Career Development Award CDA00034/2017-C to E.D. and Cross-Disciplinary Fellow LT000162/2014-C to A.J.BS), HHMI (D.B). We thank Nicolas Chiaruttini for his help with limeseg segmentation. We thank Dafne Chirivino, Vicente Planelles Herrero and Lesley McKeane for artworks. Other artworks were created with BioRender.com. We thank Xavier Morin, Lara Kruger and Buzz Baum for critical reading of the manuscript as well as Sean Munro, Gokul Upadhyayula, Lara Kruger, Karsten Kruse and Jerome Boulanger for suggestions. We thank Andrew A. Drabek and Stephen C. Blacklow for the kind gift of purified DLL4-Spytag and the U2OS-GFP-Notch cells. We are indebted to Joseph Chambers for his FLIM expertise and the CIMR light microscopy facility (Wellcome strategic award #100140) for the use of their instrument.

## Author contributions

J.L.W performed all cell experiments, except the fly panel in Fig. 5A, which A.B. performed. A.J.BS and D.B. designed the two-component 2D protein arrays and all engineered variants thereof, and A.J.BS purified the resulting proteins for use in *in vivo* assays. J.M. built the subcellular light sheet microscope, performed light sheet imaging, and wrote the deskewing and 3D deconvolution algorithm throughout this study. E.D wrote the 3D colocalization pipeline and performed image processing. J.L.W and E.D. designed the project, produced the figures and wrote the manuscript. All authors commented on the manuscript.

## Supplemental Information

Supplemental Information includes Experimental Procedures, supplementary discussion, eleven supplementary figures and ten movies.

## Abbreviations

SDCM: Spinning-Disk Confocal Microscopy
SOP: Sensory Organ Precursor

