## Supplementary data for "Induction of cortical Par complex polarity by designed proteins causes cytoskeletal symmetry breaking in unpolarized mammalian cells"

Supplementary Discussion

Experimental Procedures with Table1

Figures S1 – S11 with legends

Movie S1-S10 legends

**Supplementary discussion: Rationale for the proposed Par complex assembly paradigm**

Exactly how the three core Par complex components (Par3, Par6 and aPKC) interact with one another is a long-standing conundrum in the field. All three components are known to harbor conserved binding sites for the other components, yet the tripartite complex is not always assembled. It has often been challenging to reconcile *in vitro* data with data in cells, and to explain the different assembly states of the Par complex in different cell types and subcellular locations. In this work, through our synthetic manipulation of the system *in vivo,* we harmonize and extend previous work to provide a much fuller description of how the Par complex assembles. In this supplementary discussion, we go into more detail about how the assembly scheme depicted in Figure 3G was untangled and resolved.

It must be emphasized that all of the data on Par complex assembly detailed in this paper is highly consistent with previous work. We have confirmed the following findings:

i) Par6 and aPKC can interact, with $aPKC^{E85/R91}$ and $Par6^{K19}$ vital for this interaction (Hirano et al., 2005).
ii) Par6 and Par3 can interact, with $Par3^{G600,602}$ and $Par6^{121-257}$ vital for this interaction (Liu et al., 2020).
iii) Full length Par3 and aPKC can interact, with phosphorylation of Par3 by aPKC inhibiting this interaction (Lin et al., 2000).
iv) Clustering/oligomerization of Par3 is required for Par complex assembly (Lin et al., 2000) (Fig. S7).
v) The Par complex can assemble in clusters at cell-cell junctions (Pickett et al., 2019).

Through synthetic manipulation of individual Par complex components and mutants in 3T3 cells, we extend and reconcile these sometimes seemingly contradictory findings, which collectively suggest the molecular model depicted in Figure 3G:

1) The Par complex is mostly unassembled in single cells, with Par complex components largely cytoplasmic.
2) Clustering of any core Par complex component can induce assembly of the tripartite complex.
3) When Par3 is phosphorylated, the Par complex can still assemble via Par6, without the direct aPKC-Par3 interaction, which is therefore dispensable for assembly.
4) The aPKC kinase activity is dispensable for Par complex assembly, although its inhibition may stabilize the aPKC-Par3 interaction post-Par complex assembly.
5) Par6 exists in an autoinhibited state, with the N- and C-termini interacting to prevent binding to aPKC and Par3 respectively.
6) Clustering of any of the three components can relieve the autoinhibition of Par6 to allow binding to aPKC and Par3, but Par6 is most readily 'opened' by aPKC.

7) The role of the oligomerization domain in Par3 in Par complex assembly is solely to cluster Par3 and permit 'opening' of Par6.
These statements will be justified in the following sections.

### 1. The Par complex is natively unassembled and cytoplasmic

While Par3 and aPKC (and presumably Par6) colocalize in puncta at cell junctions (Fig. S1A-C), in naive 3T3 cells, the components are largely cytoplasmic (Fig. S1C). Colocalization cannot be reliably assessed with this diffuse protein distribution, so we sought to assess colocalization 2 minutes post induction of clustering, where assembly of lattices has long reached steady state (Fig. 1B and (Ben-Sasson et al., 2021)), and the clustered Par complex components hence appear as diffraction limited spots. At this time point, after clustering Par6, there was very little colocalization with either Par3 or aPKC (Fig. 2B,C), indicating the three components are unassembled. Furthermore, the intensity of aPKC in Par6 clusters was found to increase over time, confirming a gradual recruitment of aPKC onto these clusters, which goes against the idea of a preassembled state of the Par6/aPKC heterodimer before clustering (Fig. S5C). While endogenous Par6 could not be directly visualized, Par3 was absent from induced aPKC clusters at 2 minutes (Fig. S6C), and *vice versa* (Fig. S7D). Therefore, we conclude that the Par complex is natively unassembled in 3T3 cells.

### 2. Clustering of Par complex components can induce assembly of the tripartite complex

Synthetic clustering of either Par3, Par6 or aPKC all led to the progressive assembly of the full Par complex. Clustering Par6 led to the recruitment of aPKC followed by Par3 (Fig. 2C, Fig. S5). Clustering aPKC led to the recruitment of Par3 (Fig. S6) and Par6 (Fig. S8), and similarly, clustering of Par3 led to the recruitment of aPKC and presumably Par6 (Fig. S7).

### 3. The aPKC-Par3 interaction is insufficient and dispensable for Par complex assembly

As aforementioned, aPKC clustering induces recruitment of Par6 and Par3. This recruitment is sequential, with Par3 recruited through Par6. This is evidenced by the fact that abolishing the aPKC-Par6 interaction ($aPKC^{\Delta Par6}$) completely abolishes Par3 recruitment, irrespective of the kinase activity of aPKC (compared to $aPKC^{active\ \Delta Par6}$ and $aPKC^{dead\ \Delta Par6}$) (Fig. S6). Similarly, clustering of a short isoform of Par3 which totally lacks the aPKC binding site (100kD, noted $Par3^{\Delta aPKC}$) induced assembly of the Par complex (Fig. S7G). In this case, as we ensured binding to alternative, native Par3 isoforms was abolished through removal of the N-terminal oligomerization domain, we conclude that Par complex assembly induced by Par3 clustering must also occur through Par6. This was confirmed when we mutated the Par6 binding site on this Par3 short isoform ($Par3^{\Delta aPKC-\ \Delta Par6}$) and found that it was indeed unable to recruit aPKC (Fig. S7F). Therefore, a direct binding between aPKC and Par3 is not required for assembly of the tripartite Par complex.

However, full length Par3 (180kD) does contain an aPKC binding site so one can wonder if while the Par3/aPKC interaction is dispensable for assembly, it can regulate or potentiate it. Similarly, to $Par3^{\Delta aPKC}$, $Par3^{180kD}$ clustering was sufficient to induce Par complex assembly, with the kinetics of aPKC recruitment virtually identical between the short and full-length isoforms (Fig. S7D). This suggests that Par complex assembly induced by full length Par3 also occurs through Par6, rather than utilizing the direct aPKC-Par3 interaction. In support of this, clustering a double phosphomimetic $Par3^{180kD}$ (at positions S824 and S826, mouse equivalents of sites previously shown to inhibit binding to aPKC when phosphorylated (Lin et al., 2000)) led to Par complex assembly not significantly different to clustering the WT $Par3^{180kD}$ isoform (Fig. S7E). This confirms that the aPKC/Par3 interaction is not required for Par complex assembly.

Consistent with these findings, experiments clustering Par6 also demonstrate that Par6 is the intermediary between aPKC and Par3, rather than the aPKC-Par3 interaction driving Par complex assembly. While clustering Par6 leads to the recruitment of aPKC and the delayed recruitment of Par3 (Fig. 2C), the interaction between aPKC and Par3 is not required for this delayed recruitment of Par3. This is evidenced by Fig. 3A-E, where clustering of the N-terminus of Par6 (containing the aPKC binding site) can recruit only aPKC, and clustering of the C-terminus of Par6 (which contains the Par3 binding site) can recruit only Par3. The fact that the N-terminus of Par6 bound to aPKC cannot recruit Par3, and the C-terminus of Par6 bound to Par3 cannot recruit aPKC confirms again that the aPKC-Par3 interaction is insufficient to assemble the tripartite complex.

Nevertheless, our data are fully consistent with the fact that the phosphorylation of Par3 can prevent its direct binding to aPKC. Indeed, when we engineered a Par3 mutant that could only assemble *directly* with aPKC because it simultaneously lacks the N-terminal oligomerization domain (and therefore should not bind to alternative, native Par3 isoforms) and Par6 binding (G600,602A mutations), we found that further addition of the S824/S826 phosphomutant mutation did significantly increase its binding to aPKC (Fig. S7F, green versus yellow curves). This implies that this construct *is* able to directly recruit aPKC after clustering and thus demonstrates that, if Par3 is suitably dephosphorylated, it can promote Par complex assembly through its direct interaction with aPKC.

However, all of the evidence discussed above demonstrates that the actual route used by cells for Par complex assembly does not utilize the direct recruitment of aPKC to Par3 (or *vice versa*). In other words, it *can* happen, but it is probably not the preferred assembly route in vivo.

In conclusion, to assemble the tripartite Par complex, the interaction between aPKC and Par3 seems not required, and is natively insufficient to trigger assembly of the complex in the absence of binding to Par6. This likely explains seemingly puzzling previous observations: how could the tripartite complex assemble if the activity of aPKC towards Par3 was inhibitory for Par complex assembly, and thereby assembly would directly lead to disassembly? This could not be rationalized if the Par3/aPKC interaction was required for Par complex assembly, but in our new paradigm where it is not, the Par complex is able to assemble no matter the Par3 phosphorylation state.

It must be emphasized that this does not preclude the development of direct Par3/aPKC binding *after* Par complex assembly (via Par6). Given the proposed ability of Par6 to inhibit aPKC activity (Yamanaka et al., 2001), it is possible that, in the assembled Par complex, sufficient dephosphorylation of Par3 may occur, to permit binding between aPKC and Par3. Our data is broadly consistent with this model (as the full length Par6 better recruits Par3 than the Par6 C-terminus alone, see Fig. 3C,D), but further work will be needed to fully delineate how, if and when the aPKC-Par3 interaction can arise.

### 4. The kinase activity of clustered aPKC does not affect Par complex assembly

As clustering aPKC can trigger assembly of the tripartite Par complex, we could test the effect of kinase activating (A129E) and inhibiting (K274W) mutations, which we found to not influence Par complex assembly (Fig. S6). This is consistent with Section 3, as clustering aPKC induces Par complex assembly through Par6, which is unaffected by aPKC phosphorylation, rather than through Par3, which *is* known to be affected by aPKC phosphorylation.

One caveat here is that, when we express the kinase-dead aPKC mutant, this does not abolish all aPKC activity in the cells, as the wild-type aPKC is still expressed in the cells. Hence, it is likely that Par3 remains (at least partially) phosphorylated in these cells, and hence, the assembly route involving direct interaction between Par3 and aPKC remains unavailable. However, we can conclude that the kinase activity of aPKC *within the Par clusters* does not affect Par complex assembly.

### 5. Par6 exists in an autoinhibited state

Section 3 demonstrates that Par6 is the crucial component in Par complex assembly, as it is the intermediary between Par3 and aPKC. But then why doesn't Par6 bind to either Par3 or aPKC endogenously, when not clustered (Fig. 2B-C and Fig. S1)?

Clustering of WT Par6 leads to the rapid recruitment of aPKC, and delayed recruitment of Par3, and when Par6 cannot bind to aPKC (Par6A$^{\Delta aPKC}$ mutant), Par3 is not recruited to Par6 (Fig. 2B-D). As the aPKC-Par3 interaction cannot initiate Par complex assembly (section 3), aPKC binding to WT Par6 must change Par6 in some way, so as to allow it to bind to Par3. As discussed in section 4, this is unlikely to be due to a change in aPKC kinase activity. But how then is Par6 altered? The key insight came from clustering the C-terminus of Par6. The C-terminus contains the Par3 binding site, and lacks the aPKC binding site. It is therefore similar to the Par6A$^{\Delta aPKC}$ mutant, which also contains a Par3 binding site and lacks binding to aPKC. However, the two constructs behaved markedly differently: only the C-terminus alone of Par6 was able to recruit Par3 upon clustering (Fig. 3C,D). This demonstrates that the N-terminus of Par6 can inhibit the binding of the C-terminus to Par3. It also suggests that aPKC binding to the N-terminus of Par6 can relieve this inhibition.

Consistent with Par6 existing in a directly autoinhibited state are the experiments shown in Fig. 3F and Fig. S9, which demonstrate both in cells (with mitochondria relocalization) and *in vitro* that the N- and C-termini of Par6 can bind to one another. Further evidence comes from Fig 3C,E, where it is clear that the N-terminus of Par6 can recruit aPKC to a greater extent than the WT Par6, consistent with the C-terminus of WT Par6 partially inhibiting the aPKC binding of Par6.

We therefore conclude that Par6 natively exists in an autoinhibited state, where the N- and C-termini interact to restrict access to the aPKC and Par3 binding sites. Upon clustering of Par complex components, likely through an increase in local concentration and an enhanced avidity between components, this autoinhibition can be overcome by binding of aPKC or Par3 to Par6, which permits the binding of the remaining Par complex component to the free terminus of Par6.

### 6. Par6 autoinhibition can be relieved by Par3 or aPKC, but aPKC is more effective

In the previous section we discussed the evidence for Par6 existing in an autoinhibited state, which likely explains why the Par complex is natively unassembled in 3T3 cells. As aforementioned, clustering of any Par complex component is sufficient to induce Par complex assembly, likely by increasing the local concentration of Par complex components to a point where the affinity/avidity boost allows Par6 to be 'opened'.

Three lines of evidence suggest that aPKC is a more proficient opener of Par6 than Par3. Firstly, when Par6 itself is clustered, autoinhibition can only be relieved by aPKC (as Par6A$^{\Delta aPKC}$ cannot be 'opened' by Par3, Fig. 2C,D), at least at the local density of Par6 we achieve with our synthetic clustering. Secondly, when Par complex assembly is induced by aPKC or Par3 clustering, it is evident that aPKC more robustly assembles the Par complex than Par3 does (~42% colocalization vs ~32%, Fig. S6I vs Fig. S7D). Further, when clustering the N- or C-termini fragments of Par6, the N-terminus better recruits aPKC compared to the C-terminus recruiting Par3 (Fig. 3C-E), again consistent with the Par6-aPKC interaction being higher affinity than the Par6-Par3 interaction.

**7. The main role of Par3 oligomerization/condensation in Par complex assembly is to cluster Par3**

Previous work has demonstrated that Par3 oligomerizes *in vivo* through self-assembly of its N-terminal domain (Benton and St. Johnston, 2003; Mizuno et al., 2003). Importantly, this oligomerization has been shown to be required for assembly of the Par complex *in vivo* in flies. Our data suggest that this oligomerization likely contributes to Par complex assembly by increasing the local concentration of Par3, allowing the 'opening' of Par6. When we clustered a Par3 mutant lacking this N-terminal domain at exactly the same density as the WT, its ability to 'open' Par6 and induce Par complex assembly was comparable to that of the WT Par3 (Fig. S7G).

More recently, it has been proposed that Par3 forms condensates by phase separation (Liu et al., 2020). Combined with previous results, our data suggests that it is the high local concentration/density, rather than the specific biology and properties of biomolecular condensates (fluidity, high-density homotypic interactions, heterogenous molecular orientation, size heterogeneity, 3D geometry) that triggers Par complex assembly, as our crystalline, single layered, ordered 2D assembly could recapitulate this phenomenon (Fig. S7). Furthermore, the artificial clustering of an N-terminal truncation of Par3, which lacks the oligomerization domain reported to trigger phase separation, leads to quantitative Par complex assembly to the same extent as the full length (Fig. S7G). This suggests that the clustering density achieved by our arrays is above the threshold for maximal Par complex assembly (our arrays fix the distance between Par complex subunit at ~8 nm. For comparison, this would correspond to a concentration of ~0.55 mM to reach the same intra-subunit distance in 3D). In other words, Par complex assembly, in our system, does not depend on the combination of synthetic Par complex clustering *and* phase separation of endogenous Par3 into these clusters, but solely on the local density of Par3 achieved by our clustering method. We conclude that, whether or not Par3 phase separates in a given system, Par complex assembly is driven by an increased local density of Par3, rather than by biophysical properties of a biomolecular condensate.

**Towards a unified model of Par complex assembly**

From the conclusions detailed above, we propose a unified model for core Par complex assembly (Fig. 3G). This presents the kinetically favoured assembly route as a function of the subunit clustered.

Top row: Par6 exists natively in an unbound, autoinhibited state (sections 1 and 5). Upon clustering, aPKC can bind and 'open' Par6 (section 3 and 6), allowing the recruitment of Par3 to the C-terminus of Par6, which may be subsequently stabilized in the complex through the interaction with aPKC (section 3).

Middle row: aPKC exists natively in an unbound state (section 1), but upon clustering, independent of its kinase activity, aPKC can bind to Par6 and relieve its autoinhibition (section 6). This allows the recruitment of Par3 to the 'opened' C-terminus of Par6, and the subsequent stabilizing of Par3 into the complex if Par3 undergoes dephosphorylation (section 3).

Bottom row: Par3 natively exists in an unbound state (section 1), but, upon clustering, it can partially 'open' Par6. This allows the subsequent recruitment of aPKC into the complex, which, if Par3 is dephosphorylated, can be further stabilized through its interaction with Par3.

**Relevance of these findings to *in vivo* Par complex biology**

The work presented here highlights how synthetic biology can further our understanding of cell biology by interrogating biological systems in an orthogonal way. The key finding that the rate-limiting step in the assembly of the Par complex is the 'opening' of Par6 potentially resolves multiple discrepancies between different systems. In some systems (Benton and St. Johnston, 2003; Lin et al., 2000; Ohno, 2001), the Par complex is natively assembled, and this assembly depends on the N-terminal oligomerization of Par3. This contrasts with other systems (Morais-de-Sá et al., 2010; Soriano et al., 2016), and our findings described in section 1, where, in 3T3 cells, despite the expression of Par3 containing an N-terminal oligomerization domain, the Par complex is not assembled. We propose that, in 3T3 cells, the local concentration of Par3, even despite oligomerization/condensation, is insufficient to induce 'opening' of Par6, unlike in Drosophila, where oligomerized (but not monomeric) Par3 can 'open' Par6, likely due to a higher relative local concentration. The potential stabilizing effect of the Par3-aPKC interaction could amplify the difference between these systems, as, once the threshold for Par6 'opening' has been met, the assembled Par complex could be further stabilized. In line with Par6 'opening' depending on the local concentration of Par complex components, several studies where Par complex components have been overexpressed have demonstrated that this is sufficient to induce Par complex assembly (Kono et al., 2019; Liu et al., 2020).

It is interesting to note that aPKC is seemingly more efficient at ‘opening’ Par6 than Par3, when it is Par3 that is known, *in vivo*, to oligomerize. It will be interesting to assess what relevance this finding has *in vivo*, and whether cases exist where Par complex assembly is initiated *in vivo* through clustering of aPKC.

Finally, it has been suggested that, in some systems, Par6 and aPKC form a stable subcomplex (Chen and Zhang, 2013). Why then, if aPKC is bound to and has ‘opened’ Par6, does the triple complex not constitutively form in all these cases? Relevant to this question is the observation that the clustered C-terminus of Par6 alone only modestly colocalizes with Par3, compared to clustering of the full length Par6 (and associated aPKC). Likely, the off-rate of Par3 from the C-terminus of Par6 is high, unless stabilized by the interaction with aPKC. This stabilising interaction, which is known to be regulated by the kinase activity of aPKC and the phosphorylation state of Par3, likely determines whether a Par6-aPKC subcomplex stably binds Par3, or binds/unbinds Par3 reversibly. This will further be regulated by the concentration of components known to assemble Par complex components into different complexes (such as Cdc42), which will alter the local concentration of ‘free’ Par complex components competent to assemble the Par complex. While we did not detect colocalization between Par complex components and Cdc42 in our system (data not shown), this could affect Par complex assembly in other systems.

### Experimental Procedures

#### Protein expression and purification.

A(d), B(c)-GFP, B(c)-Spy Catcher, Spytag-DLL4 and GFP proteins were expressed and purified as described (Ben-Sasson et al., 2021). SpyTag-DLL4 was purified as described (Watson et al., 2021). B(c)-SpyCatcher: SpyTag-DLL4, referred to as B(c)-SC:ST-DLL4, was generated by mixing the two proteins in a 2:1 ratio overnight at 4$^{o}$C.

#### Cell Culture

Flp-In NiH/3T3 cells (Invitrogen) were cultured in DMEM (Gibco) supplemented with 10% Donor Bovine Serum (Gibco) and Pen/Strep 100 units/ml (Gibco) at 37°C with 5% $CO_2$. Cells were transfected with Lipofectamine 3000 (Invitrogen). Stable transfectants at the same genomic locus were obtained according to the manufacturer's instructions by homologous recombination at the FRT site were selected using 100 µg/mL Hygromycin B Gold (Invivogen).

U2OS cells (ATCC, HTB-96) were cultured in DMEM (Gibco) supplemented with 10% fetal bovine serum (Gibco) and 1% Pen/Strep at 37$^{o}$C with 5% $CO_2$ and also transfected with Lipofectamine 3000. U2OS cells expressing FLAG-Notch1-EGFP chimeric receptors were grown as described previously (Malecki et al., 2006).

#### Plasmids

All the Open Reading Frames (ORFs) cloned by PCR for this study were flanked by FseI and AscI sites for convenient shuttling between compatible plasmids. The version of GFP used throughout this study is "superfolder" GFP (sfGFP), referred to as GFP for convenience.

Murine versions of Par3 (180kDa isoform) Par6B, Par6G and aPKCι (referred to as aPKC for convenience) were cloned from murine cDNA. Par6A was cloned as a synthetic gene. Variants of these proteins were cloned using a quick-change protocol (Qiagen), to generate murine versions of established mutants/truncations, or new variants based on established protein domain structures. A summary of all these variants is provided in Table1 below for convenience. These constructs included Par3$^{\Delta N}$ (corresponding to AA 83-1319 from Par3 lacking the N-terminal oligomerization domain); Par3$^{\Delta aPKC}$ (shorter 100kDa Par3 isoform lacking the aPKC binding site, AA 1-740 + AA sequence ESGT). Note that while this truncation likely has further differences than just lacking aPKC binding, in this study, we investigated it solely in the context of its lack of aPKC binding, and hence refer to it as

Par3$^{\Delta aPKC}$ for clarity); Par3$^{\Delta Par6}$ (Par3 Par6-binding mutant, G600,602A ref.(Liu et al., 2020)); Par3 phosopho mutant (S824,826A, noted SASA, murine equivalent of S827,829A ref.(Lin et al., 2000)); Par3 phosopho mimetic mutant (S824,826D, noted SDSD); aPKC Kinase active (A129E, noted aPKC$^{active}$, murine equivalent of A120E mutant, ref.(Lim et al., 1999)); aPKC Kinase dead (K274W, noted aPKC$^{dead}$, characterized in ref.(Spitaler et al., 2000)); aPKC Par6-binding mutant (E85A/E91A, noted aPKC$^{\Delta Par6}$, characterized in ref.(Hirano et al., 2005)); Par6A aPKC-binding mutant (K19A, noted Par6A$^{\Delta aPKC}$ characterized in ref.(Hirano et al., 2005)); Par6A N-terminus (Par6A$^{1\text{-}121}$); Par6 C-terminus (Par6A$^{121\text{-}346}$) and further sub-truncations (Par6A$^{121\text{-}257}$, Par6A$^{247\text{-}346}$ Par6A$^{121\text{-}346,169LGF:AAA}$). Jupiter-iRFP670, a variant of the microtubule marker Jupiter-GFP (Karpova et al., 2006) where the GFP has been replaced with iRFP670 (Shcherbakova and Verkhusha, 2013) was synthesized by IDT.

All ORFs were cloned into a pCDNA5/FRT/V5-His vector (Life technologies) for homologous recombination into the FRT site (Flip In system). This vector has been modified to be compatible with the MXS chaining system (Sladitschek et al., 2015) to allow polycistronic expression of different constructs from the same genomic locus. ORFs were expressed under the control of the EF1a promoter, or alternatively, for Doxycycline-inducible expression, the EF1a promoter was replaced by a Tet promoter, the MXS cassette CMV::rtTA3 bGHpA was ligated into the plasmid. ORFs were tagged at the N-terminus by GFP, mCherry, or the first 34 residues of the Mas70p protein (Mito Tag), shown to efficiently relocalize proteins to mitochondria in mammalian cells (Kessels and Qualmann, 2002) followed by a HA tag (referred to as Mito-HA).

Our transmembrane nanobody construct consists of an N-terminal signal peptide from the *Drosophila* Echinoid protein, followed by $(His)_6$-PC tandem affinity tags, a nanobody against GFP (Kirchhofer et al., 2010) (termed GBP for GFP Binding Peptide), a TEV cleavage site, the transmembrane domain from the *Drosophila* Echinoid protein, the VSV-G export sequence and a second copy of the GBP. This construct is referred to as GBP-TM-GBP. For direct fusion of proteins to the transmembrane domain (Fig. S2), the internal GBP was replaced by the protein of interest. The clustering dynamics of a variant of this construct without the internal GBP has been previously extensively characterized (Ben-Sasson et al., 2021).

Most plasmids contained the EF1a Jupiter-iRFP670 cassette in order to image microtubules, but we verified that the (very dim) signals of this probe did not affect our immunofluorescence measurements in the far-red channel (Fig. S4D,E).

For pull down assays in bacterial lysate, Par6A fragments were cloned into a modified pGEX vector to express a protein of interest downstream of the Gluthathione S transferase (GST) purification tag followed by TEV and 3C cleavage sequences. Similarly, Par6A fragments were cloned into a modified pET vector expressing the protein of interest downstream of $(His)_6$ and Protein C (PC) purification tags.

| Name in this study | Construct details | Comment |
|---|---|---|
| Par6A | Par6A | Par6A wild type |
| Par6B | Par6B | Par6B wild type |
| Par6G | Par6G | Par6G wild type |
| Par6A$^{\Delta aPKC}$ | Par6A$^{K19A}$ | Par6A mutant lacking direct aPKC-binding |
| Par6A$^{Nter}$ | Par6$^{1\text{-}121}$ | Par6A N-terminus (PB1 domain) |
| Par6$^{Cter}$ | Par6A$^{121\text{-}346}$ | Par6A C-terminus (CRIB/PDZ and disordered domains) |
| Par6A$^{247\text{-}346}$ | Par6A$^{247\text{-}346}$ | Par6A C-terminal disordered domain |
| Par6A$^{121\text{-}346AAA}$ | PAR6A$^{121\text{-}346}$ with LGF$^{169\text{-}171}$ mutated to Alanine | Par6A C-terminus (CRIB/PDZ and disordered domains, with PDZ motif mutated) |
| aPKC | aPKCι | aPKCι wild type |
| aPKC$^{\Delta Par6}$ | aPKCι$^{E85A/R91A}$ | aPKC mutant lacking Par6-binding |
| aPKC$^{dead}$ | aPKCι$^{K274W}$ | aPKC mutant kinase dead |
| aPKC$^{active}$ | aPKCι$^{A129E}$ | aPKC mutant kinase active |
| aPKC$^{dead\ \Delta Par6}$ | aPKCι$^{K274W/E85A/R91A}$ | aPKC mutant kinase dead + lacking direct Par6-binding |
| aPKC$^{active\ \Delta Par6}$ | aPKCι$^{A129E/E85A/R91A}$ | aPKC mutant kinase active + lacking direct Par6-binding |
| Par3 | Par3$^{180kD}$ | Par3 wild type full length |
| Par3$^{\Delta aPKC}$ | Par3$^{100kD}$ | Par3 isoform lacking direct aPKC binding |
| Par3$^{\Delta Par6}$ | Par3$^{G600A/G602A}$ | Par3 full length mutant lacking direct Par6-binding |
| Par3$^{\Delta N}$ | Par3$^{83\text{-}1319}$ | Par3 deleted for the N-terminal oligomerization domain |
| Par3$^{SA}$ | Par3$^{S824A/S826A}$ | Par3 full length, phosphomutant on aPKC sites (more aPKC binding) |
| Par3$^{SD}$ | Par3$^{S824D/S826D}$ | Par3 full length phosphomimetic on aPKC sites (less aPKC binding) |
| Par3$^{\Delta aPKC\text{-}\ \Delta N}$ | Par3$^{100kD\ \Delta 1\text{-}83}$ | Par3 isoform lacking direct aPKC binding and N-terminal oligomerization domain |
| Par3$^{SA\text{-}\Delta Par6\text{-}\ \Delta N}$ | Par3$^{83\text{-}1319\ S824A/S826A\ G600A/G602A}$ | Par3 full length lacking direct Par6 binding and N-terminal oligomerization domain combined with phosphomutant on aPKC sites (more aPKC binding) |
| Par3$^{SA\text{-}\ \Delta N}$ | Par3$^{83\text{-}1319\ S824A/S826A}$ | Par3 full length lacking N-terminal oligomerization domain combined with phosphomutant on aPKC sites (more aPKC binding) |
| Par3$^{\Delta aPKC\text{-}\ \Delta Par6\text{-}\ \Delta N}$ | Par3$^{100kD\ G600/602A\ \Delta 1\text{-}83}$ | Par3 isoform lacking direct aPKC binding, N-terminal oligomerization domain and direct Par6-binding |
| Par3$^{\Delta aPKC\text{-}\ \Delta Par6}$ | Par3$^{100kD\ G600/602A}$ | Par3 isoform lacking direct aPKC binding + mutation lacking direct Par6-binding |

**Table1. Isoforms, truncations and mutant Par complex subunits used in this study.**

**In vitro binding between Par6 domains**

Plasmids expressing (or not) fragments of Par6 were transformed into BL21 bacteria and grown in 2XTY at 37°C, before induction with 1 mM IPTG at an OD280 of 0.8, and expression overnight at 18°C. Bacteria were then rinsed with PBS, before lysis on ice in 20 mM HEPES, 100 mM Potassium Acetate, 5 % glycerol, 1 % Triton X-100, 1X cOmplete protease inhibitors (Roche), 10 µg/ml DNaseI, 5 mM $MgCl_2$, 1 mg/ml lysozyme and 1 mM DTT for 10 minutes. Cells were then centrifuged at 20,000 x *g* for 20 minutes at 4°C. Lysates from the different expressions were then mixed, and incubated with GST-4B resin for 1 hour at 4C. The resin was subsequently washed thrice with 20 mM HEPES, 100 mM potassium acetate, 5 % glycerol, before boiling in LDS loading buffer and processing for western blot.

**SDS-PAGE and Western blot**

SDS-PAGE was performed using NuPAGE 4-12% Bis-Tris gels (Life Technologies) according to the manufacturer's instructions. Gels were transferred on nitrocellulose membranes using an iBLOT (Life Technologies) according to the manufacturer’s instructions. Following Ponceau staining (Sigma), membranes were washed in TBS and blocked in TBS enriched with 5% milk powder for 20 min at RT. Primary antibodies were incubated overnight at 4°C using a solution made of 1 µg/ml antibody in TBS supplemented with 1 mM $CaCl_2$, 0.2% BSA and 0.02% Thymerosal. The membranes were then incubated using fluorescent antibodies (1/500 dilution in TBS supplemented with 1mM $CaCl_2$ and 5% milk powder), then revealed using a fluorescent scanner (Typhoon).

**Antibodies**

All antibodies were used at 1 µg/mL for both Western Blot and immunofluorescence. Atto-647N-labelled anti-α K40 acetylated tubulin (C3B9, HPA Cultures) antibody was prepared as previously described (Derivery et al., 2015). Mouse monoclonal antibody HPC4 (against the PC tag) was from Roche. Rabbit polyclonal antibody against Par3 was from Merck (#07-330). Mouse monoclonal antibody against aPKC was from Santa Cruz (sc-17837). Rabbit polyclonal anti-GST antibody was from Abcam (#ab19256). Rat monoclonal anti-HA antibody (clone 3F10) was from Roche. Alexa-555 and Alexa-A647-conjugated anti-rabbit secondary antibodies used were purchased from Thermo (A32732, A21246). Alexa-647-conjugated anti-mouse secondary antibody was also purchased from Thermo (A-11018). Donkey Cy3-conjugated anti-rat antibodies were purchased from Jackson ImmunoResearch.

**Array assembly on cells and live cell imaging**

Clustering the transmembrane construct was achieved through assembly of our two-component hexagonal arrays, essentially as described previously (Ben-Sasson et al., 2021). These two components are named A(d) and B(c)-GFP, where B(c)-GFP binds to the transmembrane segment (via the anti GFP nanobody), and A(d) clusters B into a hexagonal array of fixed dimensions. Briefly, cells stably expressing the transmembrane construct were trypsinized and spread on Fibronectin-coated (50µg/ml in PBS, 30 minutes at RT) imaging dishes (World Precision Instruments, FD35 or FD3510) for 1 hour. Cells were then incubated for 1 minute with 0.5 µM B(c)-GFP in growth medium then washed once with medium, before the addition of 0.5 µM A(d) in medium for 10 minutes (or less, if clustering was to be imaged at an earlier time point). Cells were then washed with medium to remove free array components.

When working with cells coexpressing GBP-TM-GBP and a GFP fusions, some extracellular GBP could be quenched with GFP-fusion protein in the medium due, for example, to cell death. To avoid this potential issue, cells are always treated with a quick acid wash prior to GBP-TM-GBP clustering. Specifically, cells were briefly washed thrice with 0.1 M glycine, 150 mM NaCl, pH 3.0 to remove any bound extracellular GFP, before washing with medium. Importantly, we verified that this acid washing step did not affect the clustering efficiency nor our colocalization results (Fig. S4B,C).

To image cells through division, cells were synchronized in mitosis with 30 nM nocodazole for 6-12 hours at 37°C after clustering. Cells were then gently washed twice with imaging medium, before imaging at 37°C in L15 medium (Gibco), 10% Donor Bovine Serum (Gibco) enriched with 20 mM Hepes (Gibco).

For sub-cellular light sheet imaging, 3T3 cells expressing GBP-TM-mScarlet were spread onto 5mm glass coverslips coated with fibronectin as above. Cells were then incubated with 0.5µM B(c)-GFP as above, followed by A(d) and imaging in Hepes-enriched L15 medium.

For the cap formation from Notch receptors (Fig. S3E,F), U2OS expressing Notch receptors tagged externally with GFP were incubated with 250 nM B(c)-SC:ST-DLL4 for 5 minutes, and then with A(d)-GFP for a further 5 minutes, both in culture medium. Arrays were further grown by seven cycles of sequential incubations with B(c)-SC:ST-DLL4 then A(d), each of 1 min. Cells were then incubated with 1 µM nocodazole overnight, and imaged in medium containing nocodazole.

**Immunofluorescence**

For fixed cell imaging of the central spindle, cells were prepared in the same way as for division assays, except that, after the nocodazole block, cells were washed with DMEM medium and incubated at 37°C/5% CO2 for 30 minutes, to allow cells to reach anaphase. Cells were then fixed with 4% paraformaldehyde (PFA) in PBS for 20 minutes at room temperature. Cells were then washed in PBS and subsequently permeabilized with 0.1% Triton X-100 for 5 minutes, before washing and incubation with 1% bovine serum albumin (BSA, Fisher, BP1605) for 10 minutes. Cells were then stained with Atto-647N-labelled anti-α K40 acetylated tubulin antibodies, before imaging in PBS.

For fixed cell imaging of Par complex assembly after clustering, cells at the indicated time post-clustering, were fixed as above. Cells were stained with anti aPKC and/or anti Par3 antibodies for 1 hour at RT in PBS + 0.1% BSA. After washing thrice, cells were stained with secondary antibodies for 1 hour in PBS + 0.1% BSA. Cells were then imaged in PBS. Note that while most stable cell lines plasmids contained an EF1a Jupiter-iRFP670 cassette in order to image microtubules, we verified that the dim signals of this probe did not affect our immunofluorescence measurements in the far-red channel (Fig. S4D,E). In fact, Jupiter-IRFP670 is a weak microtubule marker and its signal can only really be sporadically detected at the central spindle in anaphase (Fig. 4).

For fixed cell imaging of mitochondrial relocalization, NiH/3T3 Flp-In cells were transiently transfected with the indicated plasmids for 24 hours. Cells were then trypsinized and spread on Fibronectin-coated dishes for 1 hour. Cells were then fixed and stained as above, with an anti-HA antibody, and imaged in PBS.

**Flow cytometry**

To measure the density of active GBP-TM-fusions at the surface of cells as a function of the expression level of each construct (Fig. S2c-e), stable 3T3 cells expressing GBP-TM-fusions under Doxycycline-inducible promoter were treated with varying doses of Doxycycline for 24 h, then cells were incubated with 1 µM purified GFP in serum/HEPES-supplemented L-15 medium for 1 min at RT, then washed in PBS-1 mM EDTA and trypsinized and resuspended in serum/HEPES-supplemented L-15 medium. GFP-fluorescence per cell was then measured by Flow cytometry in an iCyt Eclipse instrument (Sony) using a 488 nm laser. Data analysis was performed using the supplier’s software package.

**Fly notum immunofluorescence.**

Fly handling was done according to standard procedures. To generate the desired genotypes and remove balancers, all experiments were performed on F1 of crosses at 25°C. Transgenes used in this study included *UAS-mRFP-Pon*$^{LD}$ (ref. (Emery et al., 2005)) and *Neur>Gal4* (ref. (Bellaiche et al., 2001)). For immunofluorescence of dividing SOPS, larvae were then shifted to 16°C until puparium and were shifted to 25°C 16h prior to dissection.

Dissected fly nota were fixed according to a method designed to preserve the microtubule cytoskeleton (Bell Jr. and Safiejko-Mroczka, 1995). Briefly, nota were first incubated in Hank's balanced salt solution (Gibco) enriched with 1mM DSP (Pierce) for 10 min at RT followed by a 10 min incubation in MTSB (microtubule stabilization buffer: 0.1M PIPES, 1mM EGTA, 4 % PEG 8000, pH 6.9) enriched with 1mM DSP, then finally in MTSB enriched with 4% PFA (Electron Microscopy Science). Nota were then permeabilized in MTSB enriched with 4% PFA and 0.2% Triton X100 then processed for immunofluorescence using Atto-647N-labelled anti-α K40 acetylated tubulin (C3B9, HPA Cultures) antibody as described (Derivery et al., 2015) and mounted in Prolong Gold antifade reagent (Molecular Probes). Imaging was performed on the Spinning Disk confocal microscopes described below.

Detailed genotype Fig. 5a,c: *w*$^{1118}$ *; ;Neur>Gal4, UAS>mRFP-Pon*$^{LD}$ */ +* (25°C)

**Fluorescence Lifetime Imaging Microscopy (FLIM)**

3T3 cells stably expressing GBP-TM-VSVG-mScarlet were spread on fibronectin-coated glass-bottom dishes, and the transmembrane construct was then clustered with B(c)-GFP and A(d) as described above, before mitotic stalling was induced with 30 nM nocodazole for 12 h. The Flipper-TR probe (Spirochrome, see ref(Colom et al., 2018)) was then added (2 µM final in L15-20 mM HEPES medium) and Flipper-TR fluorescence lifetime imaging was performed on a setup comprising a Zeiss LSM710 stand, a 63X NA 1.4 oil objective, a Zeiss 710 confocal scanner head and Time-Correlated Single Photon Counting (TCSPC) hardware from Picoquant. A 470 nm pulsed laser (Picoquant), operating at 40 MHz was used to excite the probe, and detection was performed on a gated PMA hybrid 40 detector (Picoquant) behind a 600/50 bandpass filter (Semrock). SymPhotime 2.0 software (Picoquant) was used for data analysis. Flipper-TR fluorescence lifetime was fitted to a dual exponential model. The intensity-weighted average lifetime in a Regions of Interest (ROI) encompassing either the array-containing membrane (assessed by GFP signal), or regions of the membrane devoid of arrays was then measured, followed by averaging over several cells.

**Microscopy**

All confocal imaging was performed using a custom spinning disk confocal microscope composed of a Nikon Ti stand equipped with perfect focus, a fast piezo z-stage (ASI) and a Plan Apochromat lambda 100X NA 1.45 objective (immunofluorescence experiments) or a Plan Apochromat lambda 60X NA 1.4. The confocal imaging arm is composed of a Yokogawa CSU-X1 spinning disk head and a Photometrics 95B back-illuminated sCMOS camera operated in global shutter mode and synchronized with the spinning disk rotation. Excitation was performed using 488 nm (150 mW OBIS LX), 561 nm (100 mW OBIS LS) and 637 nm (140 mW OBIS LX) lasers fibered within a Cairn laser launch. To minimize bleedthrough, single band emission filters were used (Chroma 525/50 for Alexa 488/GFP; Chroma 595/50 for mScarlet/Alexa 555/mRFP/mCherry and Chroma 655LP for Alexa647/ATTO647N) and acquisition of each channel was performed sequentially using a fast filter wheel in each arm (Cairn Optospin). To enable fast acquisition, the entire setup is synchronized at the hardware level by a Zynq-7020 Field Programmable Gate Array (FPGA) stand-alone card (National Instrument sbRIO 9637) running custom code. In particular, fast z-stacks are obtained by synchronizing the motion of the piezo z-stage during the readout time of the cameras. Sample temperature was maintained at 37°C for microtubule dynamics using a heating enclosure (MicroscopeHeaters.com, Brighton, UK). Acquisition was controlled by Metamorph software.

Sub-cellular light sheet imaging (Fig. S2B and movie S1) was performed on a home-built field synthesis (Chang et al., 2019) light sheet microscope featuring a 0.7 NA excitation lens (54-10-7, Special Optics) and a 1.1 NA detection lens (CFI75 Apo 25XC W MRD77220, Nikon). Illumination light was provided from 405 nm, 488 nm, 561 nm and 638 nm lasers (LBX-405-100-CSB-PPA, LBX-488-100-CSB-PPA and LBX-638-100-CSB-PPA, Oxxius or OBIS 561 nm LS 100 mW, Coherent, respectively). Fluorescence was collected by the detection objective and directed through an f = 500 mm doublet onto the camera (Pco Edge 4.2, PCO, giving a sample pixel pitch of 104 nm), through appropriate bandpass filters (Chroma). The sample was coarsely positioned in XY using a stack of piezo-driven translation stages (8525, Newport) and was moved through a fixed sheet during imaging using a fast-stepping piezo drive (P-621.1CD, PI). The sample was mounted on a 5 mm diameter circular cover slip (AGL46R5-1, Agar Scientific) held in a stainless steel 'spoon'. The sample was submerged in L-15 medium supplemented with HEPES held in a gold-plated copper bath, which was resistively heated at 37°C. We noted that the imaging performance of the detection objective was highly sensitive to the exact setting of the correction collar, which was manually iteratively optimized before every imaging session.

### Image processing

Unless stated otherwise, images were processed using Fiji (Schindelin et al., 2012) and Matlab 2020b (Mathworks) using custom codes. Figures were assembled in Adobe Illustrator 2021. Movies were edited in Adobe Premiere 2021.

Spatial drift during acquisition was corrected using a custom GPU-accelerated registration code based on cross correlation between successive frames. Drift was measured on one channel and applied to all the channels in multichannel acquisitions.

For representation purposes, all confocal images of Par clusters in colocalization figures (Figs. 1-3 and Data Figs. S1,4-8) were processed with a Wavelet "à trous" filter (custom GPU-accelerated MATLAB port of a code originally developed by Fabrice Cordeliere for the "Improve Kymo" ImageJ plugin (Zala et al., 2013)) and the raw image was averaged with the filtered one to generate the figure panel. Note that the colocalization analysis (see below) was performed on the raw 3D stacks.

#### 1) Subcellular light-sheet data processing

Image volumes obtained by sub-cellular light sheet microscopy were computationally deskewed using the home-made software suite lsfm_tools implementing a linear interpolation (https://github.com/jdmanton/lsfm_tools). CudaDecon (https://github.com/scopetools/cudaDecon) was used to deconvolve data using the Richardson-Lucy algorithm with a theoretical PSF (Born and Wolf model) generated with the ImageJ plugin PSF generator (Kirshner et al., 2013). Deskewed volumes were visualized via maximum intensity projection.

#### 2) 3D reconstruction

For 3D reconstruction (Fig. 1, Fig. S3D, movies S1,S3), confocal z-stack of cells (Δz=200 nm) were acquired, and deconvolved using the Richardson-Lucy algorithm and a theoretical PSF in the program suite lsfm_tools described above. Cell surface was then automatically segmented in 3D using the Fiji plugin LimeSeg developed by Machado and colleagues (Machado et al., 2019), then 3D rendering was performed using Amira software.

#### 3) Colocalization

To automatically measure the colocalization between clusters and endogenous proteins in fixed samples, we used an object-based method where two objects are considered colocalized if the distance between their fluorescence centroid is below a certain threshold $r_{ref}$ (Bolte and Cordelieres, 2006). Particles were automatically detected in multichannel confocal z-stacks (Δz=200 nm) using 3D

Gaussian PSF fitting developed by Aguet and colleagues (Aguet et al., 2016) (https://github.com/francois-a/llsmtools). This also provided the integrated intensity of each spot considering the local background. We excluded from the analysis all arrays localized at the ventral side of the cell to avoid potential artefacts due to interactions of the arrays with the coverslip, or to differential membrane tension between the ventral and dorsal sides of the cell. To achieve this, the coverslip was automatically detected, and all arrays found within 3 planes of the coverslip (600 nm) were automatically excluded. This also provided the integrated intensity of each spot considering the local background.

Once this automated detection has been performed in all channels, the distance $d_{AB}$ between all particles in 3D in the two channels ($A$ and $B$) are computed and compared to a reference distance $r_{ref}$. If $d_{AB} < r_{ref}$, the particles detected in the two channels are deemed to colocalize.

To set $r_{ref}$ we followed the method implemented by Cordelières and Bolte in the ImageJ plugin JACop 2.0 (Bolte and Cordelieres, 2006), and calculated $r_{ref}$ for the 3D situation using the following equations:

$$\Phi = \arccos\frac{(x_B - x_A)}{\sqrt{(x_B - x_A)^2 + (y_B - y_A)^2}} \text{ and } \Theta = \arccos\frac{(z_B - z_A)}{\sqrt{(x_B - x_A)^2 + (y_B - y_A)^2 + (z_B - z_A)^2}}$$

$$r_{ref} = \sqrt{\left(resol_{xy} \sin\Theta \cos\Phi\right)^2 + \left(resol_{xy} \sin\Theta \sin\Phi\right)^2 + (resol_z \cos\Theta\,)^2}$$

Here, $x_A, y_A, z_A$ and $x_B, y_B, z_B$ are the 3D coordinates of particles in channel $A$ and $B$, respectively, and $resol_{xy}$ and $resol_z$ correspond to the lateral and axial resolutions of the microscope, respectively. For our analysis, we measured $resol_z = 0.498\,\mu m$ and $resol_{xy} = 0.293\,\mu m$ using 0.2 µm TetraSpeck beads from Invitrogen.

Once all the particles have been detected and their colocalization state addressed (i.e. $d_{AB} < r_{ref}$), we measured the percentage of colocalization as the fraction of the total signal contained in particles that do colocalize, namely:

$$\%\ of\ colocalization = \frac{\sum colocalizing\ particles}{\sum total\ particles} * 100$$

This measurement was then averaged between cells and compared between stable cell lines and time points. For each graph, a sample where only GFP is clustered (rather than GFP-Par6A for instance) is plotted to serve as an internal control to show the accuracy of the method (any colocalization with GFP alone is assumed to reflect non-specific colocalization due to the random localization and density of spots in both channels).

4) Central spindle asymmetry measurements

Central spindle asymmetry was measured by computing the difference of microtubule density (assessed by acetyl-tubulin immunostaining) between the two sides of the central spindle, rather than the absolute amounts of tubulin. This is because we previously established that what matters for polarized trafficking of signalling endosomes is the ratio of tubulin densities between the two sides of the spindle, rather than the absolute amount of tubulin (Derivery et al., 2015). Briefly, we first projected z-stacks containing the entire central spindle (6 µm depth, $\Delta z = 0.2$ µm) using max-intensity projection. We focused on cells fixed shortly before abscission as at this stage, the Ac-tubulin signal at the central spindle has the characteristic hourglass shape of late mitotic spindles. We can thus use this characteristic shape as a registration cue to register spindles from different images and average them, as we previously established (Derivery et al., 2015).

The intensity of the tubulin staining was then measured along central spindle length (x-axis) upon signal integration over the spindle width (y-axis, parallel to the division plane) within a region of interest (ROI) centered on the central spindle. This measurement thus conceptually resembles a linescan along the length of the spindle (x-axis), but where not a line, but a rectangular ROI is considered (ROI dimensions: 10 µm on the x-axis and spindle width on the y-axis). For this reason, we previously named it "pseudo-linescan" method (Derivery et al., 2015). The signal intensity over the x-axis determined this way displays two peaks: one in pIIa, one in pIIb, see Fig. 5C for an example. This reflects the facts that most markers are excluded (at least in part) from the core of the central spindle. These "pseudolinescans" were then averaged between different cells to yield the panels presented in Figs.5C,F.

We then measured the value of each peak and subtracted the local background (average background was determined from 5 pixels adjacent to the spindle). Central spindle asymmetry was computed as the normalized enrichment of the density of the marker in the anterior side according to:

$$\Delta = \frac{\text{Peak intensity anterior} - \text{Peak intensity posterior}}{\text{Peak intensity anterior} + \text{Peak intensity posterior}}$$

Note that $\Delta$ is symmetrical when anterior and posterior cells are inverted and that $-1 \leq \Delta \leq 1$. Antero-posterior polarity was provided by the $Pon^{LD}$ signal for fly SOPs, or by the localization of the artificial cap for 3T3 cells (by convention, the cap-containing cell is the posterior cell as Par3/Par6/aPKC are posterior Par proteins). This value was then averaged per cell to yield Fig. 5G.

**Statistics**

Unless stated otherwise, measurements are given as mean ± SEM. Statistical analyses were performed using GraphPad Prism 8 with an alpha of 0.05. Normality of variables was verified with Kolmogorov-Smirnov tests. Homoscedasticity of variables was always verified when conducting parametric tests. Post-hoc tests and their respective p-values are indicated in their respective figure legends.

**Supplementary Figures**

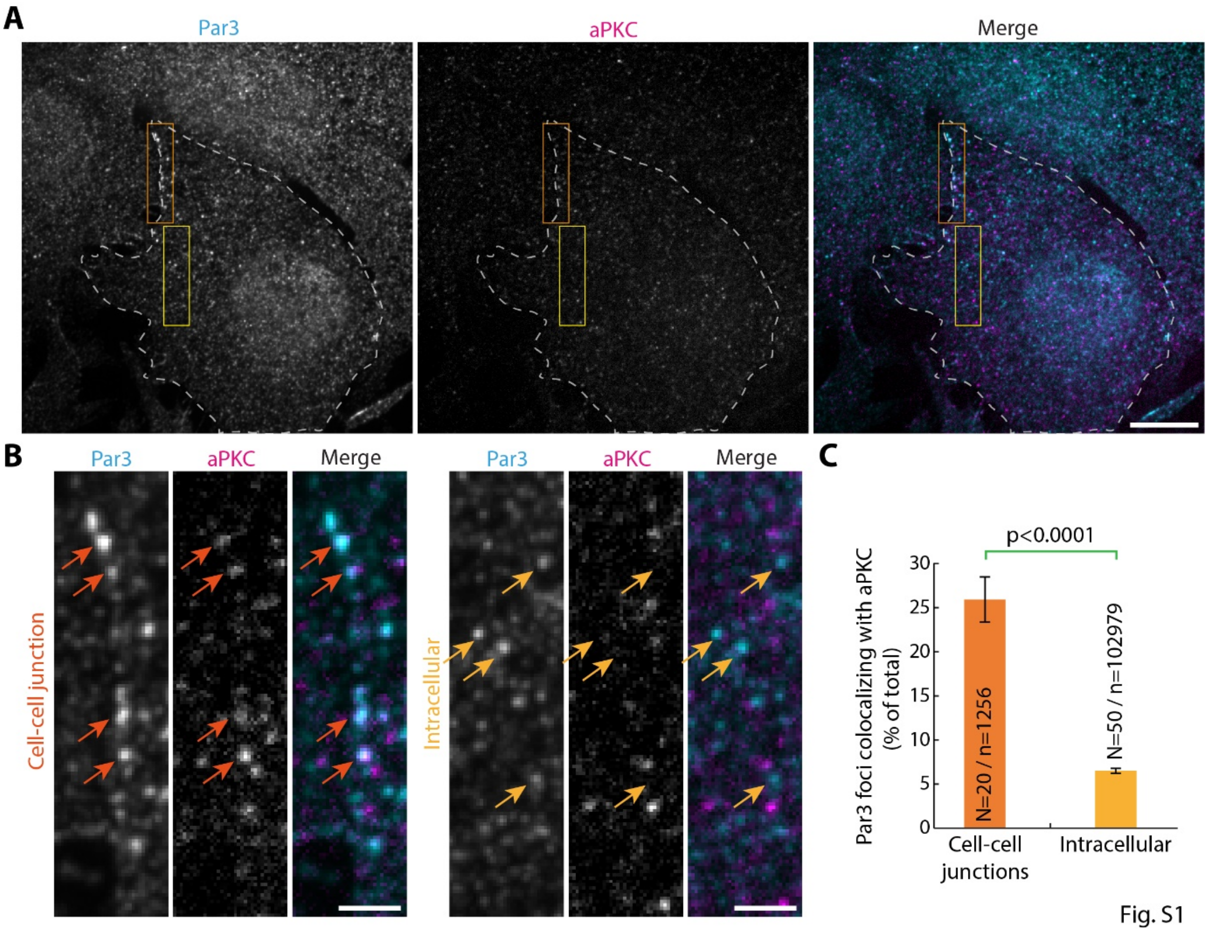

**Figure S1. The Par complex assembles at cell-cell junctions**

(**A**) 3T3 cells spread on fibronectin-coated coverslips were immunostained for endogenous Par3 and aPKC and imaged by SDCM. Images correspond to maximum-intensity projection of 7 planes (Δz=1.4µm total). Dashed white line: cell contours. (**B**) High magnification of the image presented in (**A**) in a cell-cell junction region (left panel, corresponds to orange rectangle in **A**) or in an intracellular region (right panel, corresponds to yellow rectangle in **A**). Note the colocalization between Par3 clusters and aPKC at cell-cell junctions (orange arrows), in contrast to the rest of the cell (yellow arrows). (**C**) Mean (+/-SEM) percentage of colocalization between Par3 clusters and aPKC per region (see also methods for details about the automated 3D object-based colocalization method developed for this paper). Statistics: Unpaired Student's t test (N: regions of interest analysed; n: total number of Par3 spots detected). Scale bars: 10 µm (**A**) ; 2 µm (**B**).

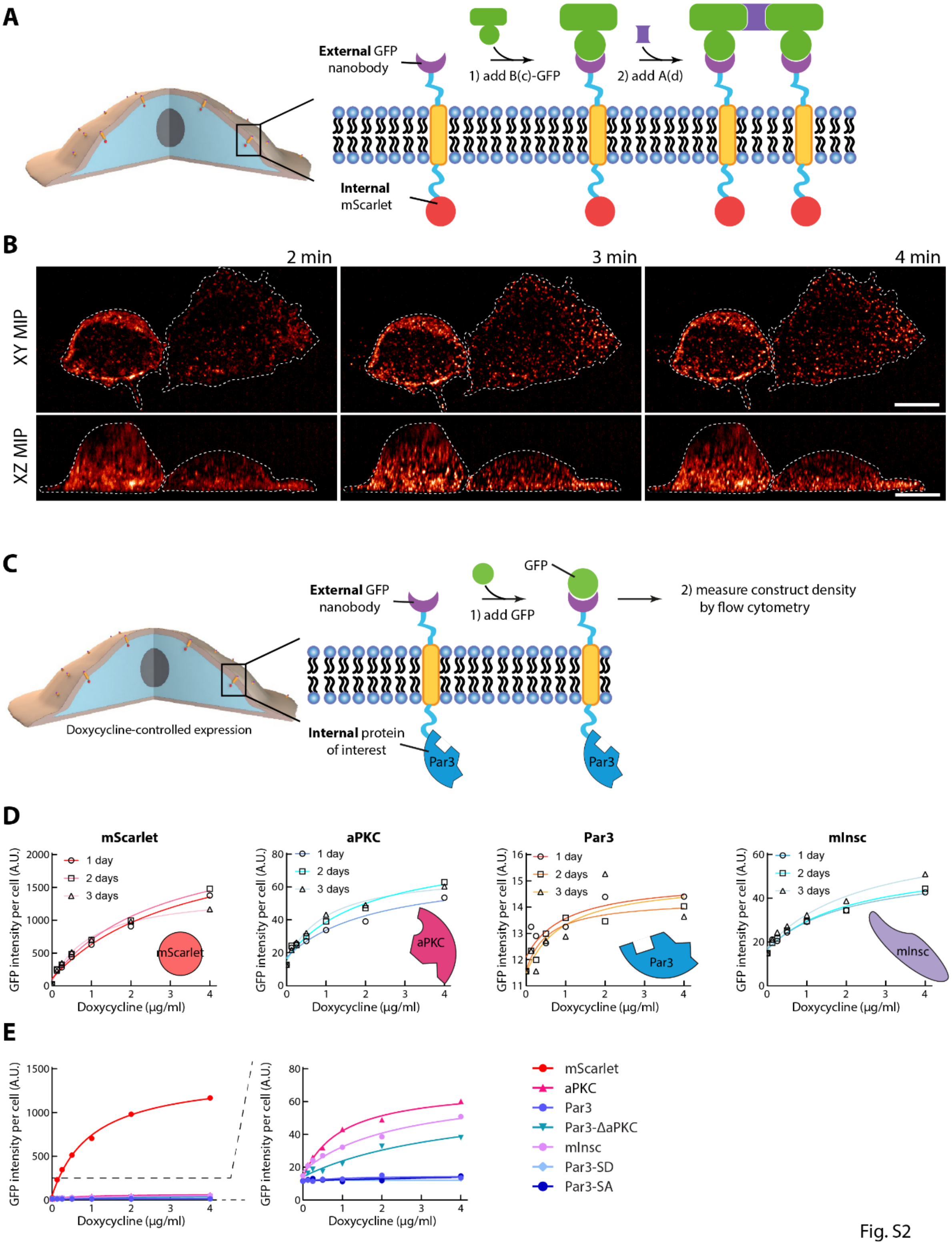
A
External GFP nanobody
1) add B(c)-GFP
2) add A(d)
Internal mScarlet
B
2 min
3 min
4 min
XY MIP
XZ MIP
C
External GFP nanobody
GFP
1) add GFP
2) measure construct density by flow cytometry
Doxycycline-controlled expression
Internal protein of interest
Par3
D
mScarlet
aPKC
Par3
mInsc
1 day
2 days
3 days
GFP intensity per cell (A.U.)
Doxycycline (µg/ml)
E
mScarlet
aPKC
Par3
Par3-ΔaPKC
mInsc
Par3-SD
Par3-SA

Fig. S2

**Figure S2. Volumetric clustering dynamics and effect of fused protein on construct density**

(**A**-**B**) 3D Dynamics of array clustering. (**A**) Principle of the experiment: 3T3 cells stably expressing GBP-TM-mScarlet protein were incubated with B(c)GFP then A(d) to induce rapid clustering. (**B**) Cells treated as in (**A**) were imaged by sub-cellular light sheet microscopy (see also movie S1). Images correspond to maximum intensity projections (from the top or the side). Note that array assembly occurs homogeneously at all points of the cell cortex. Scale bars: 10 µm. (**C**-**E**) The expression level of the transmembrane construct is heavily affected by the protein it is fused too. (**C**) Principle of the experiment: 3T3 cells expressing GBP-TM fused to a protein of interest under the control of a Doxycycline inducible promoter were treated with Doxycycline for the indicated time, before incubation with purified GFP. The GFP signal per cell is then measured by Flow cytometry as a proxy of the density of the transmembrane construct at the cell surface. (**D**) Cells expressing the indicated fusion were treated as described in (**C**). The duration of the Doxycycline treatment does not quantitatively affect the surface density of the construct. (**E**) Cells expressing the indicated fusion were treated with Doxycycline for 3 days and processed as above. The identity of the protein fused to the transmembrane construct dramatically affects the surface density of the construct. This is a concern, as we previously established that the size of 2D arrays (and thus the number of clustered receptors per array) is a direct function of the initial density of the transmembrane construct (Ben-Sasson et al., 2021). For this reason, most data in this paper relies on a bipartite system where the transmembrane construct is fused to an intracellular anti-GFP nanobody, and the protein of interest is fused to GFP (Fig. 1A and methods).

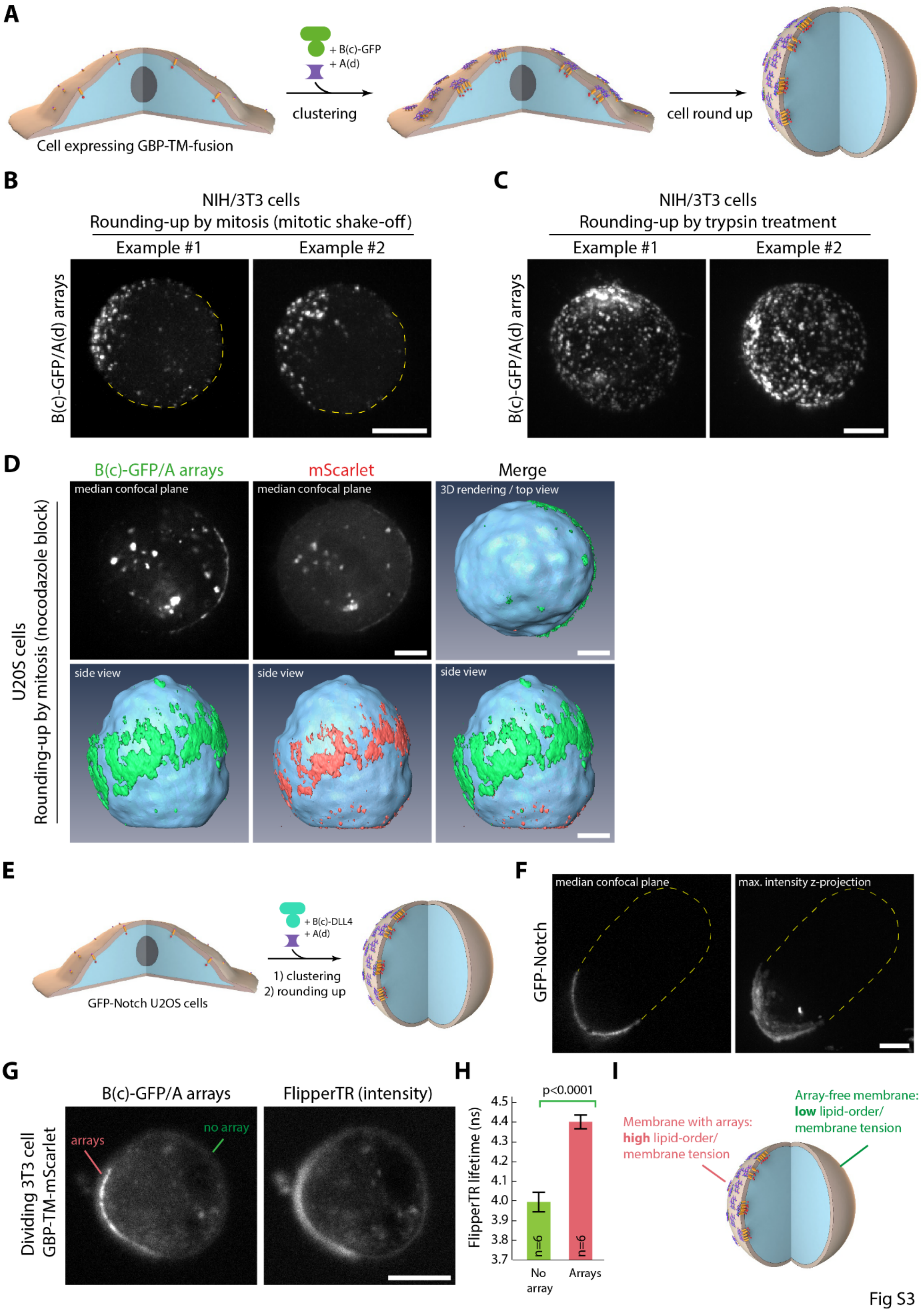
A
Cell expressing GBP-TM-fusion
+ B(c)-GFP
+ A(d)
clustering
cell round up
B
NIH/3T3 cells
Rounding-up by mitosis (mitotic shake-off)
Example #1
Example #2
B(c)-GFP/A(d) arrays
C
NIH/3T3 cells
Rounding-up by trypsin treatment
Example #1
Example #2
B(c)-GFP/A(d) arrays
D
B(c)-GFP/A arrays
mScarlet
Merge
median confocal plane
median confocal plane
3D rendering / top view
side view
side view
side view
U2OS cells
Rounding-up by mitosis (nocodazole block)
E
+ B(c)-DLL4
+ A(d)
1) clustering
2) rounding up
GFP-Notch U2OS cells
F
median confocal plane
max. intensity z-projection
GFP-Notch
G
B(c)-GFP/A arrays
FlipperTR (intensity)
arrays
no array
Dividing 3T3 cell
GBP-TM-mScarlet
H
p<0.0001
FlipperTR lifetime (ns)
4.5
4.4
4.3
4.2
4.1
4.0
3.9
3.8
3.7
n=6
n=6
No array
Arrays
I
Membrane with arrays:
high lipid-order/
membrane tension
Array-free membrane:
low lipid-order/
membrane tension

Fig S3

**Figure S3. Cap formation is a general property of 2D arrays and is not restricted to cell type or cell-interfacing modalities**

(**A**) Principle of the experiment used in this figure. Array are assembled at the surface of cells, either via a synthetic GFP/GBP link, or via an endogenous ligand/receptor pair, and then ability of arrays to coalesce into a polar cap upon cell rounding up is assessed. (**B-C**) Cell rounding-up induces cap formation, rather than specifically mitosis. Sequential incubation with B(c)GFP (1 min, 0.5 µM) and A(d) (5 min, 0.5 µM) was used to assemble arrays at the surface of 3T3 cells coexpressing GBP-TM - GBP and GFP-LGN (**B**) or expressing GBP-TM-mScarlet (**C**). The arrays were then imaged by SDCM upon rounding up by entering mitosis (**B** mitotic shake-off), or by trypsin treatment (**C**). Two representative examples of cap formation in both cases are shown. (**D**) Cap formation is not restricted to 3T3 cells. Arrays were assembled at the surface of U2OS cells transiently expressing GBP-TM-mScarlet using B(c)GFP and A as above. Cells were then stalled in mitosis with 40 nM nocodazole for 12 h and imaged by SDCM followed by 3D reconstruction and surface rendering (see methods). Note the presence of a polar cap positive for GFP and mScarlet. The high overexpression in this transient expression experiment (compared with the low-level stable expression in the rest of this study) likely explains the higher presence of intracellular GFP/mScarlet staining. (**E**-**F**) Cap formation is not restricted to artificial GBP-TM constructs. (**E**) Principle of the assay: Arrays are assembled at the surface of U2OS cells expressing GFP-Notch by sequential incubation with 0.5 µM Bc-SC:ST-DLL4 (DLL4 is a Notch ligand) and A(d)-GFP, followed by few rounds of further sequential growth (see methods). (**F**) Cells treated as in (**E**) were imaged by SDCM. Note the Notch-positive polar cap. (**G**-**I**): Array oligomerization locally modifies the biophysical properties of the membrane they are attached to. (**G**) 3T3 cells stably expressing GBP-TM-mScarlet were spread on fibronectin-coated imaging dishes, and array assembly was then triggered by sequential incubation with B(c)-GFP and A(d). Cells were then stalled in mitosis with 30 nM nocodazole for 12h. Cells were then incubated with Flipper-TR probe (2 µM final in L15-20 mM HEPES medium) and fluorescence lifetime imaging was performed in the same medium (see methods). (**H**) Intensity-weighted average lifetime of the Flipper-TR probe in region where arrays are present (segmented using GFP fluorescence) or not (mean ± SEM). A higher Flipper-TR lifetime of the probe indicates a local high-order and/or high-tension in the membrane. Statistics: paired Student's t test (p value indicated; n: number of cells analyzed). (**I**) Putative mechanism of array coalescence: Array assembly creates a localized region of higher membrane tension and/or higher lipid packing compared to the surrounding naked membrane. This creates a line tension around the domain, which will tend to induce coalescence of arrays into a cap by energy minimization. Scale bars: 5 µm.

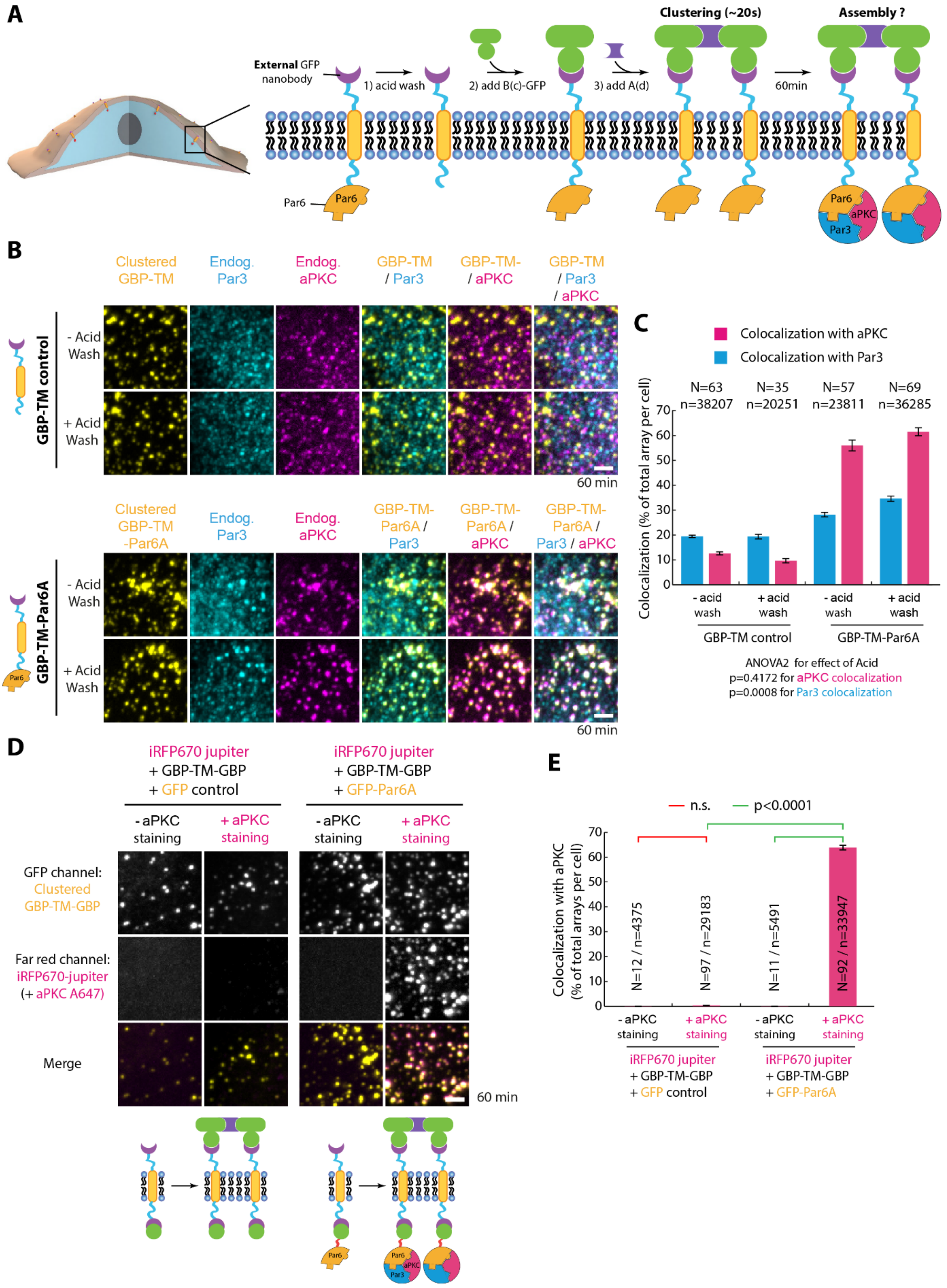
A
External GFP nanobody
1) acid wash
2) add B(c)-GFP
3) add A(d)
Clustering (~20s)
60min
Assembly ?
Par6
aPKC
Par3
B
Clustered GBP-TM
Endog. Par3
Endog. aPKC
GBP-TM / Par3
GBP-TM- / aPKC
GBP-TM / Par3 / aPKC
GBP-TM control
- Acid Wash
+ Acid Wash
60 min
Clustered GBP-TM -Par6A
GBP-TM-Par6A / Par3
GBP-TM-Par6A / aPKC
GBP-TM-Par6A / Par3 / aPKC
GBP-TM-Par6A
C
Colocalization with aPKC
Colocalization with Par3
N=63 n=38207
N=35 n=20251
N=57 n=23811
N=69 n=36285
Colocalization (% of total array per cell)
- acid wash
+ acid wash
GBP-TM control
GBP-TM-Par6A
ANOVA2 for effect of Acid
p=0.4172 for aPKC colocalization
p=0.0008 for Par3 colocalization
D
iRFP670 jupiter + GBP-TM-GBP + GFP control
iRFP670 jupiter + GBP-TM-GBP + GFP-Par6A
- aPKC staining
+ aPKC staining
GFP channel: Clustered GBP-TM-GBP
Far red channel: iRFP670-jupiter (+ aPKC A647)
Merge
60 min
E
n.s.
p<0.0001
Colocalization with aPKC (% of total arrays per cell)
N=12 / n=4375
N=97 / n=29183
N=11 / n=5491
N=92 / n=33947

Fig. S4

**Figure S4. Control experiments of the bicistronic clustering technology**

(**A-C**) The acid wash treatment to ensure that the external GBP is not bound to GFP does not affect the accuracy of the colocalization measurements with aPKC or Par3 used in this study. (**A**) Principle of the experiment. When working with cells coexpressing GBP-TM-GBP and a GFP fusions, some extracellular GBP can be quenched with GFP-fusion protein ended up in the medium due to cell death, for example. To avoid this issue, cells are always treated with a quick acid wash prior to GBP-TM-GBP clustering. To test if this acid wash affects the accuracy of our aPKC / Par3 colocalization pipeline, 3T3 cells expressing GBP-TM-Par6A (or GBP-TM control) were spread on fibronectin-coated coverslips. Cells were processed for acid-wash followed by clustering of the GBP-TM construct by incubation with B(c)GFP then A(d). In this experiment, the extracellular GBP cannot be quenched by GFP-fusions at the beginning of the experiment as the only source of GFP is exogenous B(c)GFP. Thus, measuring the colocalization between Par6 and aPKC / Par3 allows one to measure the effects of the acid wash. (**B**) Cells were processed as in (**A**), then were immunostained for endogenous aPKC and Par3 after 60 min of clustering. Imaging was performed by SDCM. Images correspond to maximum intensity projection ($\Delta z$ =3.8-4.6µm total). (**c**) Mean (+/-SEM) percentage of the 3D colocalization between GFP-Par6 (or GFP control) clusters and aPKC or Par3 per cell. Effect of the acid wash was tested using an ANOVA2 test (respective p-value indicated). Acid wash has only a negligible effect on the extent of colocalization between Par6, aPKC and Par3. (**D-E**) Weak iRFP670-Jupiter signal in the far-red channel does not affect the accuracy Alexa-647 aPKC immunostaining. (**D**) 3T3 cells expressing iRFP670-Jupiter, GBP-TM-GBP and GFP-Par6A (or GFP control) were spread on fibronectin-coated coverslips, then GBP-TM-GBP was clustered using B(c)GFP and A(d) as in Figure 1. Cells were then immunostained (or not) for endogenous aPKC using Alexa647-coupled secondary antibodies. Cells were then imaged by SDCM. Images correspond to maximum intensity projection ($\Delta z$ =3.4-4.6µm total). Dynamic range was set to be identical between images. (**E**) Mean (+/-SEM) percentage of the 3D colocalization between Par3 clusters and aPKC per cell. Statistics were performed using a Kruskal-Wallis test followed by a Dunn post-hoc test (p-value of respective tests indicated). Throughout this figure, N corresponds to the number of cells analyzed per condition, and n to the total number of GFP-positive arrays detected per condition. All images in this figure were processed with a wavelet “a trous” filter (see methods). Scale bars: 2 µm (**B, D**).

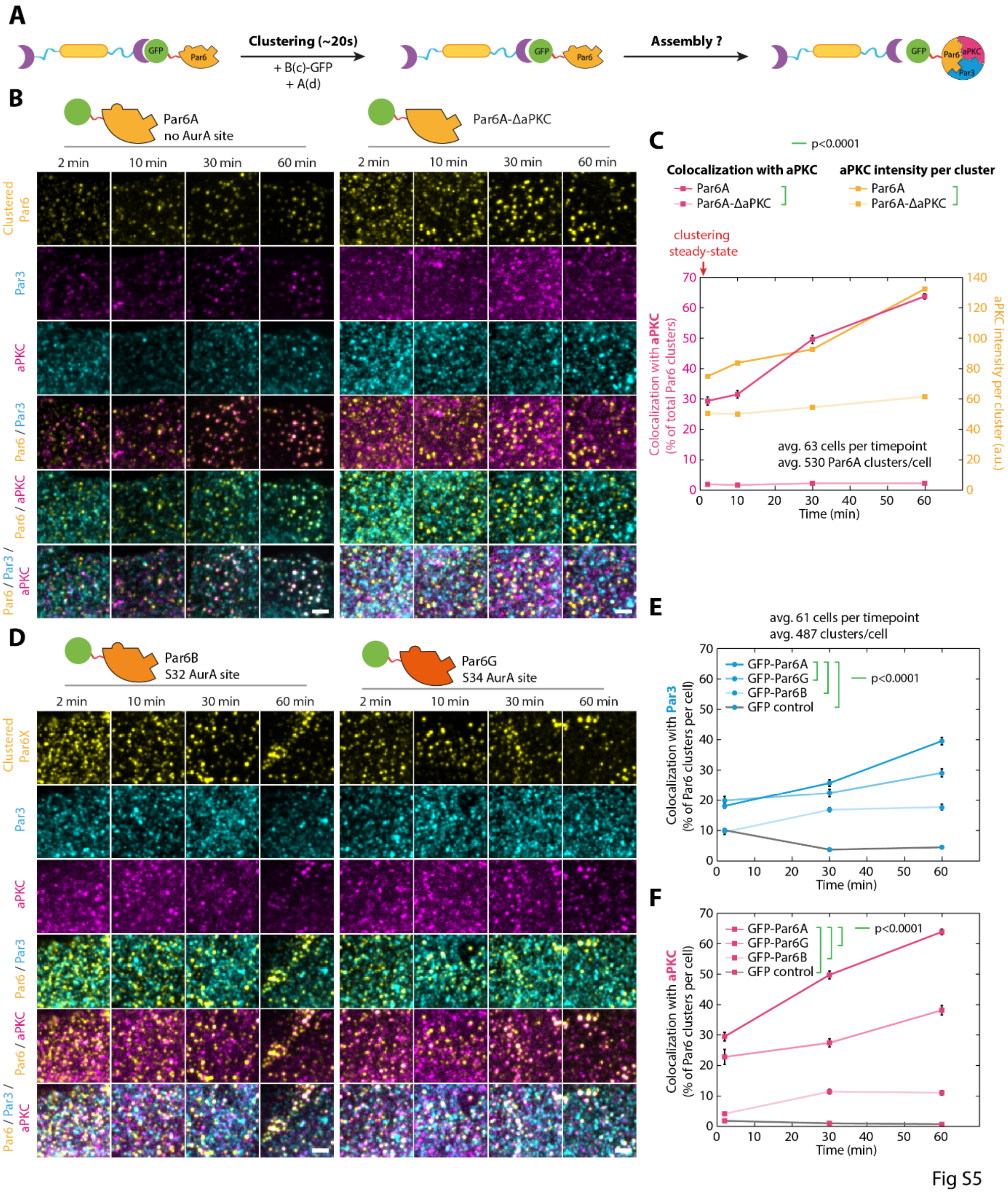

### Figure S5. Variable ability of Par6 isoforms to support Par complex assembly

(**A**) Principle of the experiment: 3T3 cells stably co-expressing GBP-TM-GBP and GFP-fused Par6 variants were incubated with B(c)GFP then A(d) to induce rapid clustering, and the recruitment of endogenous aPKC and Par3 was monitored by immunofluorescence. (**B**) Cells expressing indicated Par6 mutant were treated as presented in (**A**), and the recruitment of endogenous aPKC and Par3 was monitored by immunofluorescence. This corresponds to the split channel panels of Fig. 2. (**C**) Mean

(+/-SEM) percentage of colocalization between GFP-Par6A (or GFP-Par6A$^{\Delta aPKC}$) clusters and aPKC over time, plotted at the same time as the mean (+/-SEM) intensity of aPKC per cluster over time. Both the binary colocalization and the aPKC intensity (i.e. the number of molecules recruited) increase over time, suggesting slow recruitment of aPKC molecules onto stable Par6A clusters. Note that the mean percentage of colocalization curves are the same as the ones presented in Fig. 2C, shown here for convenience. (**D**) Kinetics of recruitment of aPKC and Par3 to clusters of indicated Par6 isoforms assessed by immunofluorescence. (**E-F**) Mean (+/-SEM) percentage of colocalization between clusters of indicated Par6 isoform and Par3 (**E**) or aPKC (**F**). Statistics: ANOVA2 using construct and timepoint as variables followed by (p-value of each test indicated). While all Par6 isoforms can assemble the Par complex (colocalization is significantly above the control), the three isoforms are markedly different in the speed at which they assemble the core Par complex upon clustering. Note that the Par6A and GFP-control curves are the same as the ones presented in Fig. 2C, shown here for convenience. All images in this figure were processed with a wavelet “a trous” filter (see methods). Scale bars: 2 µm (**B**,**D**).

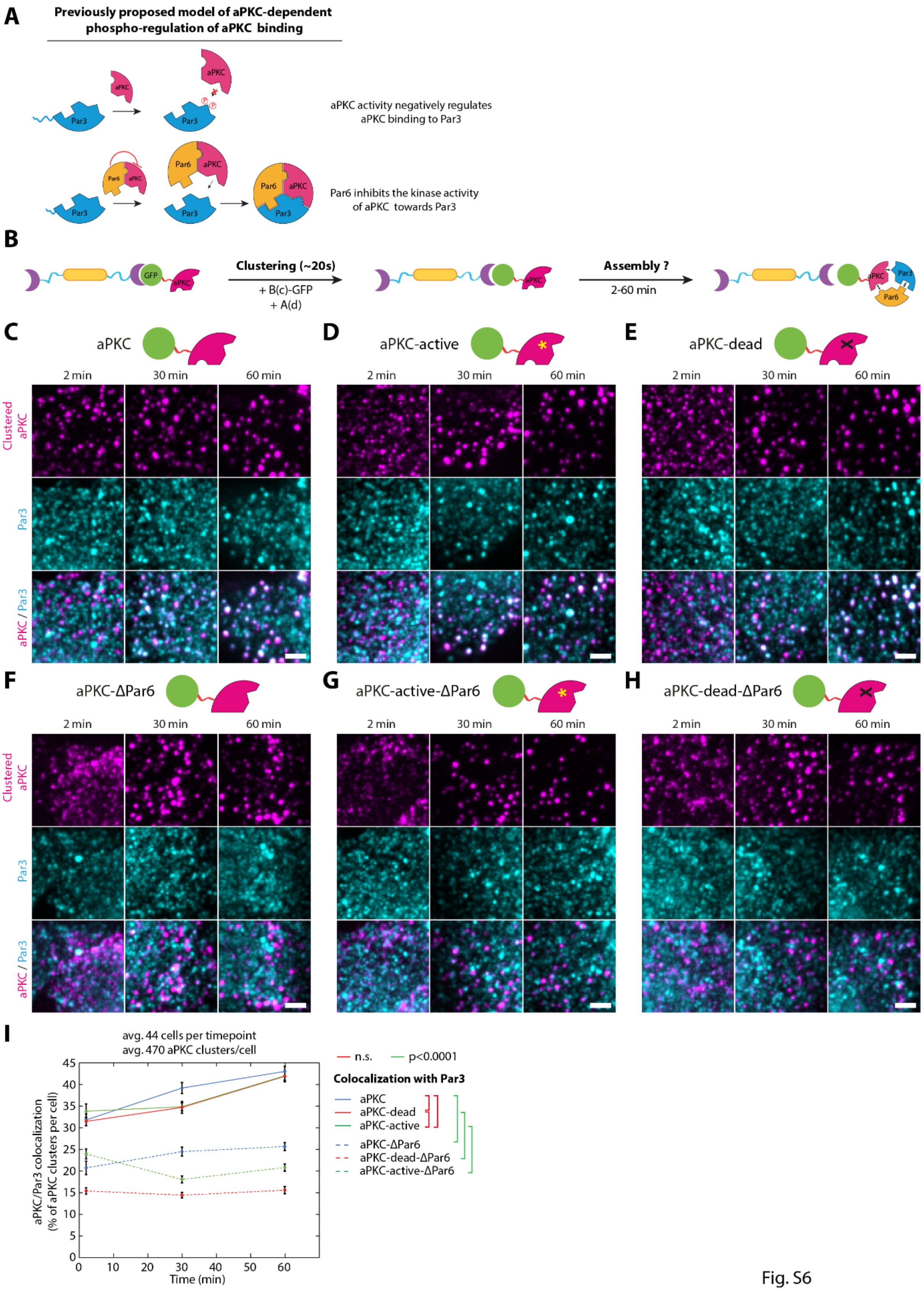
A
Previously proposed model of aPKC-dependent phospho-regulation of aPKC binding
aPKC activity negatively regulates aPKC binding to Par3
Par6 inhibits the kinase activity of aPKC towards Par3
B
Clustering (~20s)
+ B(c)-GFP
+ A(d)
Assembly ?
2-60 min
C
aPKC
D
aPKC-active
E
aPKC-dead
2 min
30 min
60 min
Clustered aPKC
Par3
aPKC / Par3
F
aPKC-ΔPar6
G
aPKC-active-ΔPar6
H
aPKC-dead-ΔPar6
I
avg. 44 cells per timepoint
avg. 470 aPKC clusters/cell
aPKC/Par3 colocalization (% of aPKC clusters per cell)
Time (min)
n.s.
p<0.0001
Colocalization with Par3
aPKC
aPKC-dead
aPKC-active
aPKC-ΔPar6
aPKC-dead-ΔPar6
aPKC-active-ΔPar6

Fig. S6

**Figure S6. aPKC clustering induces Par complex assembly independently on the kinase activity of clustered aPKC.**

(**A**) Previously proposed model of regulation of Par complex assembly via aPKC activity from the literature (see supplementary discussion). (**B**) Principle of the experiment: 3T3 cells stably expressing GBP-TM-GBP and GFP-fused aPKC mutants were incubated with B(c)GFP then A(d) to induce rapid clustering. (**C**-**H**) Cells expressing indicated aPKC mutant were treated as presented in (**B**), and the recruitment of endogenous Par3 was monitored by immunofluorescence. (**I**) Mean (+/-SEM) percentage of colocalization between clusters of indicated aPKC mutant and Par3. Statistics: ANOVA2 using construct and timepoint as variables followed by (p-value of each test indicated). The kinetic activity of the clustered aPKC has no impact on its ability to assemble the core Par complex, and Par3 recruitment to aPKC clusters likely occurs via Par6. All images in this figure were processed with a wavelet “a trous” filter. Scale bars: 2 µm.

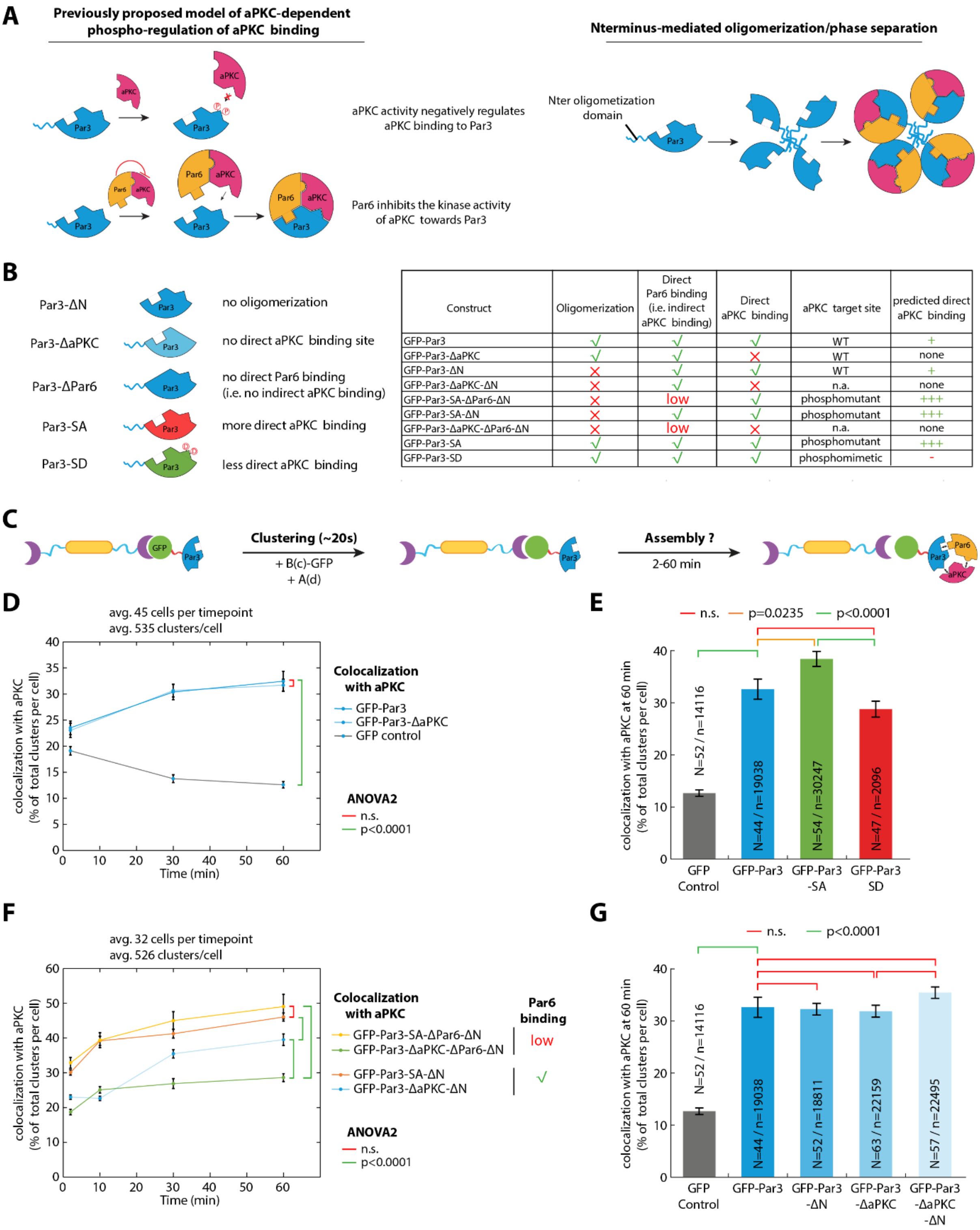

| Construct | Oligomerization | Direct Par6 binding (i.e. indirect aPKC binding) | Direct aPKC binding | aPKC target site | predicted direct aPKC binding |
|---|---|---|---|---|---|
| GFP-Par3 | √ | √ | √ | WT | + |
| GFP-Par3-ΔaPKC | √ | √ | × | WT | none |
| GFP-Par3-ΔN | × | √ | √ | WT | + |
| GFP-Par3-ΔaPKC-ΔN | × | √ | × | n.a. | none |
| GFP-Par3-SA-ΔPar6-ΔN | × | low | √ | phosphomutant | +++ |
| GFP-Par3-SA-ΔN | × | √ | √ | phosphomutant | +++ |
| GFP-Par3-ΔaPKC-ΔPar6-ΔN | × | low | × | n.a. | none |
| GFP-Par3-SA | √ | √ | √ | phosphomutant | +++ |
| GFP-Par3-SD | √ | √ | √ | phosphomimetic | - |

Fig S7

**Figure S7. Par3 clustering induces Par complex assembly without requiring direct binding to aPKC.**

(**A**) Left: Previously proposed model of regulation of Par complex assembly via aPKC activity from the literature (see supplementary discussion). Note that this panel is identical to Fig. S6A, reproduced here for convenience. Right: Proposed model of regulation of Par complex assembly via Par3 oligomerization/condensation via the N-terminal domain. (**B**) Description of the different Par3 constructs used in this figure, and their predicted effects on oligomerization, aPKC binding and Par6 binding (see also methods). (**c**) Principle of the experiment: 3T3 cells stably expressing GBP-TM-GBP and GFP-fused Par3 (or mutant thereof), were incubated with B(c)GFP then A(d) to induce rapid clustering. Then, the assembly of endogenous Par complex was followed over time by aPKC immunofluorescence and automated quantification. (**D-G**) Mean (+/-SEM) percentage of colocalization between clusters of GFP-Par3 (or indicated Par3 mutant) and aPKC, either as a time course (**D**,**F**) or as endpoint measurements (**E**,**G**). Clustering of GFP alone, which does not bind to aPKC is provided as a negative control to assess the accuracy of the method in all cases. Statistics: ANOVA2 using time and construct as variables (**D**,**F)**; p-value for effect of the construct indicated, average number of cells/array per cell per timepoint indicated) or ANOVA1 followed by a Tukey post-hoc test (**E**,**G**; p-value for effect of each post-test indicated; N: number of cells quantified; n: total number of arrays quantified per condition). See supplemental discussion for details.

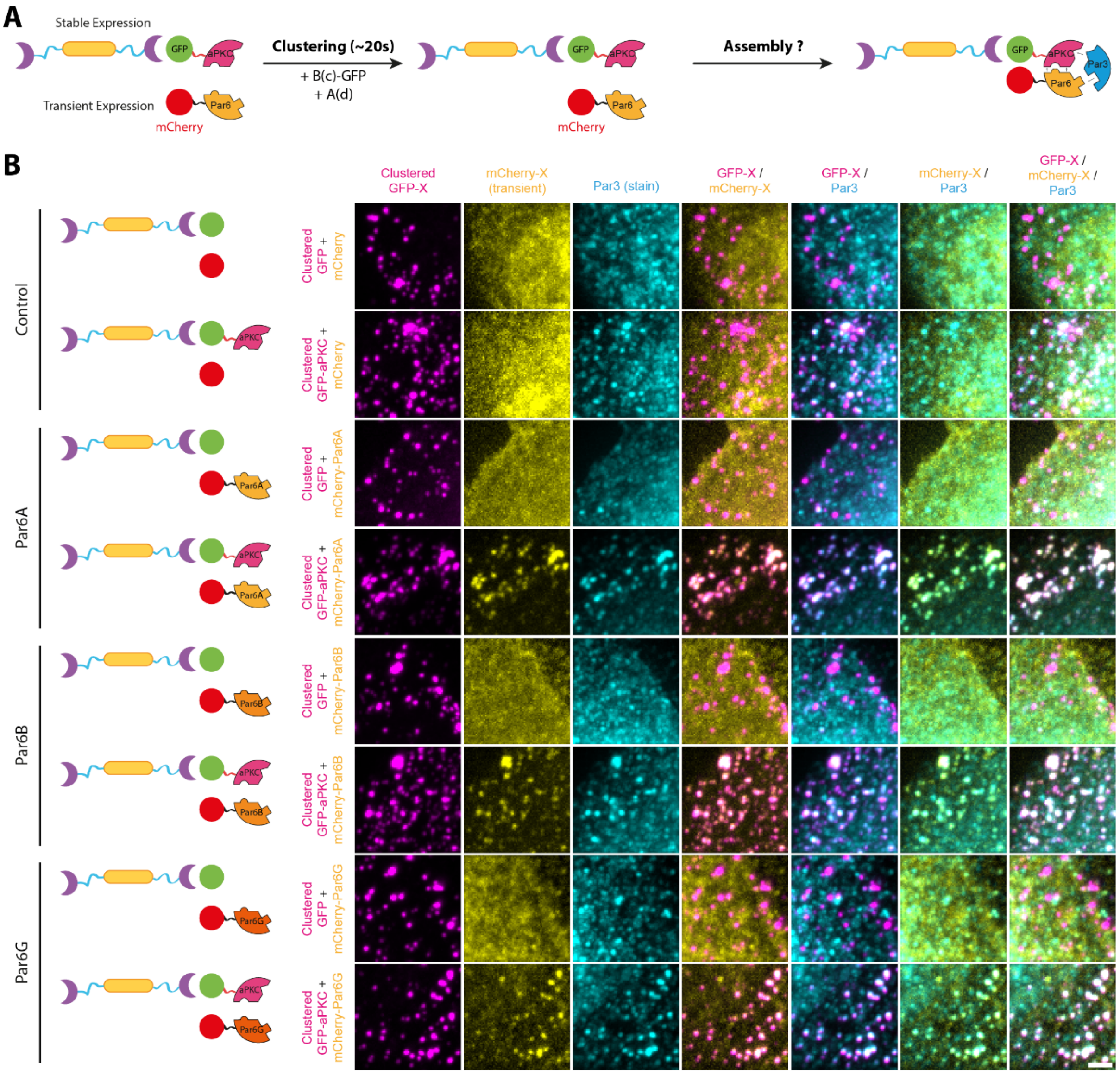

Fig S8

**Figure S8. All Par6 isoforms can be recruited onto clustered aPKC**

(**A**) Principle of the experiment: 3T3 cells stably expressing GBP-TM-GBP and GFP-fused aPKC, and transiently transfected with mCherry-Par6 isoforms were incubated with B(c)GFP then A(d) to induce rapid clustering. (**B**) Cells expressing indicated Par6 variants (or controls) were treated as presented in (**A**), and the recruitment of exogenous Par6 and endogenous Par3 to aPKC clusters was monitored by immunofluorescence. All Par6 isoforms can be recruited to aPKC clusters. All images in this figure were processed with a wavelet “a trous” filter. Scale bars: 2 µm.

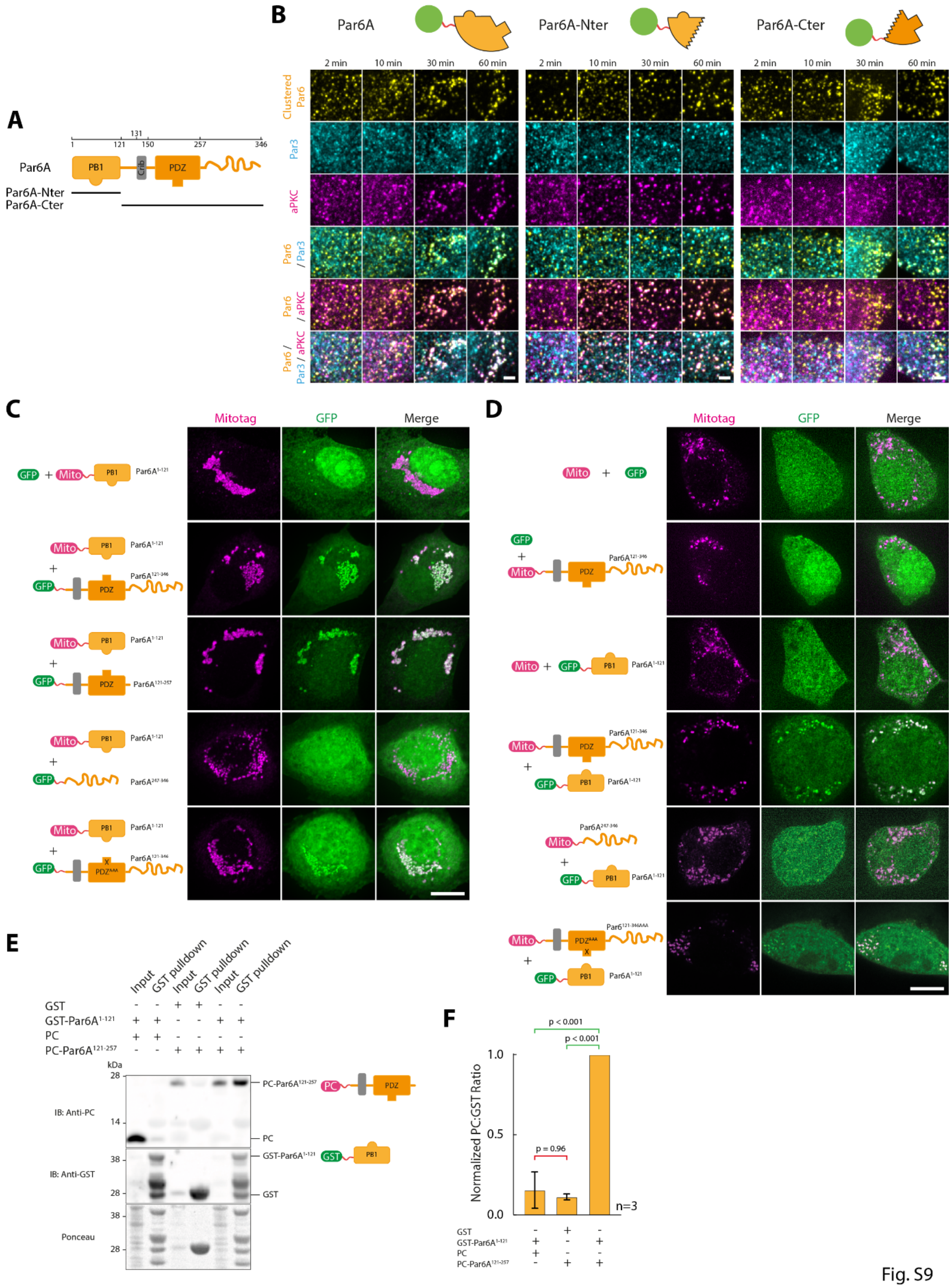
A
Par6A
PB1
Crib
PDZ
Par6A-Nter
Par6A-Cter
B
Par6A
Par6A-Nter
Par6A-Cter
2 min
10 min
30 min
60 min
Clustered Par6
Par3
aPKC
Par6 / Par3
Par6 / aPKC
Par6 / Par3 / aPKC
C
Mitotag
GFP
Merge
D
E
Input
GST pulldown
GST
PC
kDa
IB: Anti-PC
IB: Anti-GST
Ponceau
F
Normalized PC:GST Ratio
p < 0.001
p = 0.96
n=3

Fig. S9

**Figure S9. Par6 is autoinhibited**

(**A**) Domain organization of Par6A, with the N-terminal PB1 domain binding to aPKC and the C-terminal PDZ domain binding to Par3. (**B**) GFP-fused Par6A, or fragments thereof, were clustered in 3T3 cells as in Fig. 1 and the recruitment of endogenous aPKC/Par3 was monitored by immunofluorescence. Images correspond to maximum intensity projection and images were denoised with a "Wavelet a trous" filter. This corresponds to the split-channel images of the data presented in Fig. 3C. (**C**-**F**) The N-terminal domain of Par6A can bind to the C-terminus. (**C**) 3T3 cells coexpressing the indicated C-terminal fragment of Par6A tagged with GFP, and the N-terminal half of Par6 tethered to the mitochondria, or relevant controls, were assessed for GFP-recruitment to the mitochondria by SDCM. Images correspond to maximum intensity z-projection over the entire cell. Par6 $PDZ^{AAA}$ corresponds to the Alanine mutation of AAs 169_LGF_171 in full length Par6, known to abolish binding of PDZ domains. (**D**) Converse experiment as the one in (**C**), where the N-terminal fragment of Par6A is fused to GFP and coexpressed with the indicated C-terminal fragments localized to mitochondria. The N-ter and C-ter of Par6 interact but this does not go through canonical PDZ binding. (**E**,**F**) One half of Par6 can pull down the other when expressed in bacteria, suggesting a direct interaction. (**E**) GST-$Par6A^{1-121}$ and PC-$Par6A^{121-346}$ were expressed independently in bacteria, and the bacteria lysates were then mixed and incubated with glutathione beads to pull down the GST tag. The presence of PC-$Par6A^{121-346}$ was then assessed by western blot (picture representative of n=3 experiments). (**F**) Quantification of the effects seen in (**E**), suggesting specific pull down of PC-$Par6A^{121-346}$ in presence of GST-$Par6A^{1-121}$. Note that $Par6A^{1-121}$ is referred to as $Par6A^{Nter}$ in the main text, and $Par6A^{121-346}$ to $Par6A^{Cter}$. Scale bars: 2 µm (B) and 10 µm (C,D).

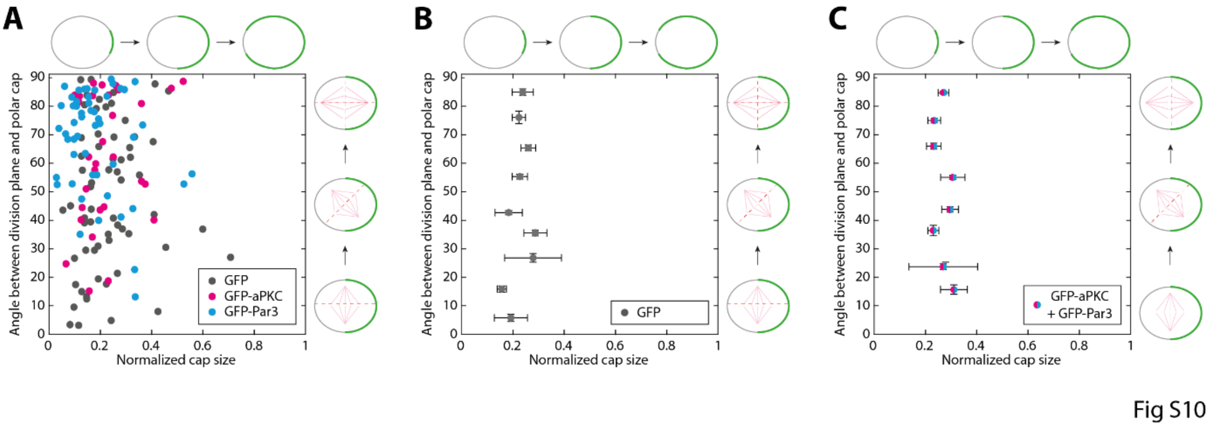

**Figure S10. Cap size has no influence on spindle orientation**

(**A**) Array assembly was triggered at the surface of 3T3 cells stably co-expressing Jupiter-iRFP670, GBP-TM-GBP as well as indicated GFP-fusions. Cells were then stalled in mitosis for >12h to form caps, and then imaged by SDCM upon release of the nocodazole block. The angle between the division plane and the array cap as a function of the protein targeted to the cap was then measured, as well as the size of the cap, and all data points were plotted on the same graph. To have a normalized measurement between cells, cap size was expressed as a fraction of the cell perimeter, and was determined in metaphase in the equatorial plane. This plot was used to generate Fig. 4G after binning. (B,C) Individual plots of the overlaid plot presented in Fig. 4G.

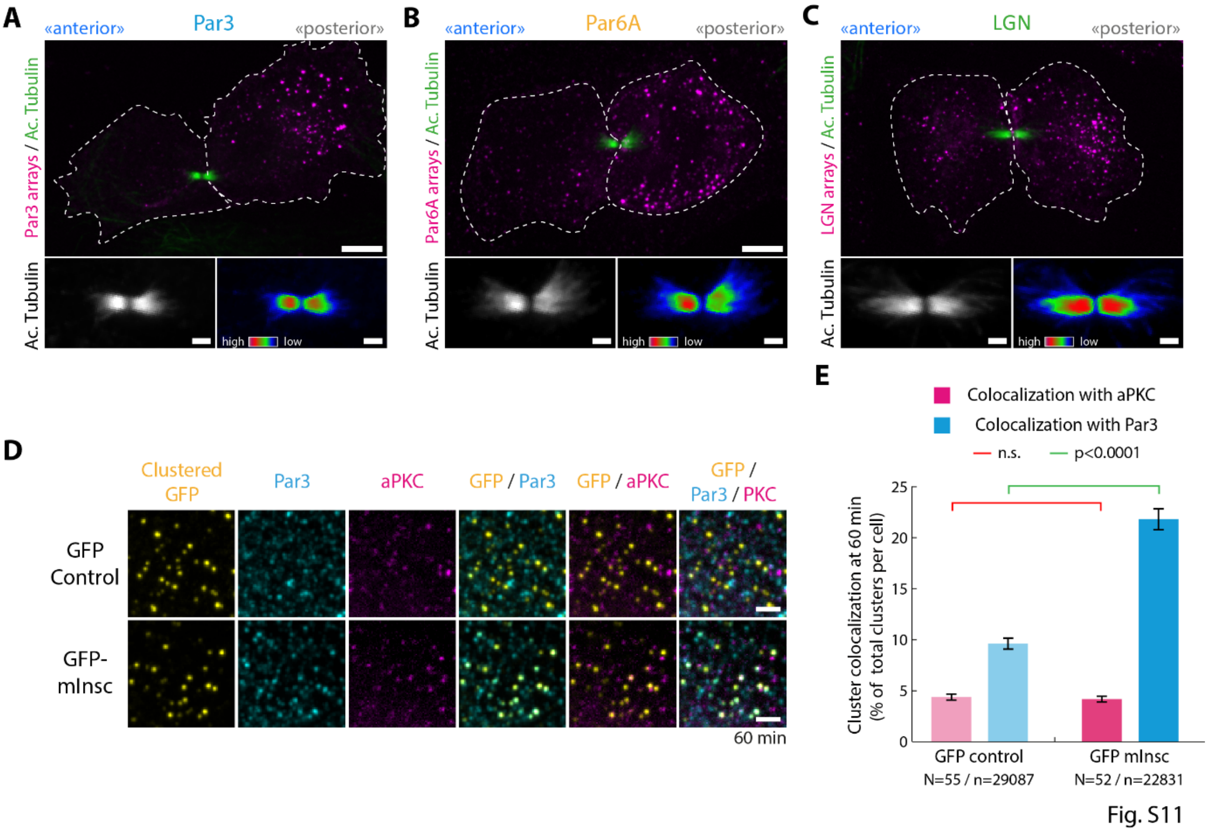

**Figure S11. An asymmetric Par complex cortex is sufficient to induce central spindle asymmetry**

(**A**-**C**) Array assembly and cap formation was triggered at the surface of 3T3 cells stably expressing GBP-TM-GBP, as well as GFP-fusions with indicated proteins. Cells were then stalled in mitosis for >12 h. Nocodazole block was then released and cells were fixed after 30 min and processed for acetyl-tubulin immunostaining and SDCM imaging (maximum intensity projection). Bottom panels: split-acetyl-tubulin channel with a grayscale (left) or Rainbow (right) lookup table. Note that in the case of Par3 (**A**) and Par6A (**B**), the side of the central spindle in the cell that did not inherit the cap ("anterior") has a higher density of microtubule than the "posterior" side, as in SOP cells (Fig. 5A) as in the aPKC case (Fig. 5B). This is not true for LGN (**C**). (**D**) Array assembly was triggered at the surface of 3T3 cells stably co-expressing Jupiter-iRFP670, GBP-TM-GBP as well as GFP-mInsc or GFP as a control. Recruitment of endogenous aPKC and Par3 was then assessed by immunofluorescence. For representation purposes, images were processed with a wavelet "a trous" filter. (**E**) Mean (+/-SEM) percentage of the 3D colocalization between GFP-mInsc (or GFP control) clusters and aPKC or Par3 per cell. Statistics: student's t-test (N: number of cells quantified; n: total number of arrays quantified per condition). While mInsc clusters do recruit Par3, they do not recruit aPKC. Scale bars: 5 µm (**A-C**, top panels) and 1 µm (**A**-D).

**Supplementary Movie legends**

**Movie S1. Volumetric imaging of array clustering**

3T3 cells stably expressing GBP-TM-mScarlet were sequentially incubated with B(c)GFP and A(d) to induce rapid clustering, and imaged by sub-cellular light sheet microscopy. Note that array assembly occurs homogeneously at all points of the cell cortex. This movie corresponds to Fig. S2A,B.

**Movie S2. Spontaneous cap formation in 3T3 cells during mitosis**

Array assembly was triggered at the surface of 3T3 cells stably expressing GFP-LGN and GBP-TM-GBP by sequential incubation with B(c)GFP (1 min, 0.5 µM) and A(d) (5 min, 0.5 µM). Cells were then stalled in mitosis for >12h with nocodazole, and then imaged by SDCM, followed by 3D reconstruction and surface rendering (see methods). Note the partitioning of arrays into an asymmetric cortical cap. This movie corresponds to Figure 1D.

**Movie S3. Spontaneous cap formation in U2OS cells during mitosis**

Arrays were assembled at the surface of U2OS cells transiently expressing GBP-TM-mScarlet by sequential incubation with B(c)GFP (1 min, 0.5 µM) and A(d) (5 min, 0.5 µM). Cells were then stalled in mitosis with 40 nM nocodazole for 12 h and imaged by SDCM followed by 3D reconstruction and surface rendering (see methods). Note the presence of a polar cap positive for GFP and mScarlet. The high overexpression in this transient expression experiment (compared with the low-level stable expression in the rest of this study) likely explains the higher presence of intracellular GFP/mScarlet signal. This movie corresponds to Fig. S3D.

**Movie S4. An asymmetric cortex of GFP does not orient mitotic spindle**

Array assembly was triggered at the surface of 3T3 cells stably expressing GFP and GBP-TM-GBP by sequential incubation with B(c)GFP (1 min, 0.5 µM) and A(d) (5 min, 0.5 µM). Cells were then stalled in mitosis for >12h with nocodazole, and then imaged by SDCM. Movie starts in metaphase. Note that the division plane is not aligned with the cap formed by the arrays, and thereby array segregation is symmetrical.

**Movie S5. An asymmetric cortex of GFP-Par3 induces spindle orientation**

Array assembly was triggered at the surface of 3T3 cells stably expressing GFP-Par3 and GBP-TM-GBP by sequential incubation with B(c)GFP (1 min, 0.5 µM) and A(d) (5 min, 0.5 µM). Cells were then stalled in mitosis for >12h with nocodazole, and then imaged by SDCM. Movie starts in metaphase. Note that the division plane is aligned with the cap formed by the GFP-Par3 arrays, and thereby array segregate preferentially into one cell.

**Movie S6. An asymmetric cortex of GFP-aPKC induces spindle orientation**

Array assembly was triggered at the surface of 3T3 cells stably expressing GFP-aPKC and GBP-TM-GBP by sequential incubation with B(c)GFP (1 min, 0.5 µM) and A(d) (5 min, 0.5 µM). Cells were then stalled in mitosis for >12h with nocodazole, and then imaged by SDCM. Movie starts in metaphase. Note that the division plane is aligned with the cap formed by the GFP-aPKC arrays, and thereby array segregate preferentially into one cell.

**Movie S7. An asymmetric cortex of GFP-Par6 induces spindle orientation**

Array assembly was triggered at the surface of 3T3 cells stably expressing GFP-Par6 and GBP-TM-GBP by sequential incubation with B(c)GFP (1 min, 0.5 µM) and A(d) (5 min, 0.5 µM). Cells were then stalled in mitosis for >12h with nocodazole, and then imaged by SDCM. Movie starts in metaphase. Note that the division plane is aligned with the cap formed by the GFP-Par6 arrays, and thereby array segregate preferentially into one cell.

**Movie S8. The kinase activity of aPKC is dispensable for spindle orientation**

Array assembly was triggered at the surface of 3T3 cells stably expressing GFP-aPKC$^{dead}$ and GBP-TM-GBP by sequential incubation with B(c)GFP (1 min, 0.5 µM) and A(d) (5 min, 0.5 µM). Cells were then stalled in mitosis for >12h with nocodazole, and then imaged by SDCM. Movie starts in metaphase. Note that the division plane is aligned with the cap formed by the GFP-aPKC$^{dead}$ arrays, and thereby array segregate preferentially into one cell.

**Movie S9. An asymmetric cortex of GFP-mInsc induces spindle orientation**

Array assembly was triggered at the surface of 3T3 cells stably expressing GFP-mInsc and GBP-TM-GBP by sequential incubation with B(c)GFP (1 min, 0.5 µM) and A(d) (5 min, 0.5 µM). Cells were then stalled in mitosis for >12h with nocodazole, and then imaged by SDCM. Movie starts in metaphase. Note that the division plane is aligned with the cap formed by the GFP-mInsc arrays, and thereby array segregate preferentially into one cell.

**Movie S10. An asymmetric cortex of GFP-LGN induces spindle orientation** Array assembly was triggered at the surface of 3T3 cells stably expressing GFP-LGN and GBP-TM-GBP by sequential incubation with B(c)GFP (1 min, 0.5 µM) and A(d) (5 min, 0.5 µM). Cells were then stalled in mitosis for >12h with nocodazole, and then imaged by SDCM. Movie starts in metaphase. Note that the division plane is aligned with the cap formed by the GFP-LGN arrays, and thereby array segregate preferentially into one cell.
